# Genome-wide CRISPRi screen identifies basigin loss as protective in cardiac hypoxia

**DOI:** 10.64898/2026.01.26.701810

**Authors:** Will R. Flanigan, Ayush D. Midha, Skyler Y. Blume, Yolanda Marti-Mateos, Mauro W. Costa, Yu Huang, Alan H. Baik, Helen Huynh, Gautam Susarla, Neal K. Bennett, Romana A. Nowak, Deepak Srivastava, Ken Nakamura, Isha H. Jain

## Abstract

Cardiac function depends on continuous oxidative metabolism, rendering cardiomyocytes highly vulnerable to oxygen deprivation. Here, we performed a genome-wide CRISPR interference (CRISPRi) screen in human iPSC-derived cardiomyocytes to identify genes that modulate survival during chronic hypoxia. This screen revealed that knockdown of basigin (BSG), a chaperone for the monocarboxylate transporters MCT1 and MCT4, confers robust protection. Canonically, hypoxic cells suppress pyruvate dehydrogenase (PDH) activity to reduce the oxidation of major fuel sources, thereby limiting TCA cycle flux, lowering oxygen consumption, and minimizing reactive oxygen species generated by an overly reduced electron transport chain (ETC). In contrast, we found that BSG inhibition reverses this response, prioritizing ATP maintenance during hypoxia and enhancing cardiomyocyte survival. Mechanistically, BSG loss restricts lactate efflux, leading to decreased PDH phosphorylation and increased glucose uptake for oxidation. Consistent with this, ETC subunits are more essential under hypoxia, highlighting cardiomyocytes’ unusual reliance on aerobic ATP production even when oxygen is limited. These findings challenge prevailing models of hypoxic adaptation by revealing cardiomyocyte-specific bioenergetic requirements and motivating future therapeutic efforts.

## INTRODUCTION

The heart is an obligate aerobic organ with one of the highest oxygen demands in the human body.^1^ Its continuous mechanical activity requires a constant oxygen supply to sustain oxidative phosphorylation and ATP generation. Even brief interruptions in oxygen delivery can compromise contractile function, while prolonged oxygen deprivation leads to irreversible injury. Consequently, hypoxia (low oxygen supply) and ischemia (low blood supply) underlie a broad spectrum of cardiovascular diseases,^2^ including ischemic heart disease, heart failure, and cyanotic congenital heart disease.^3^ At present, we lack effective treatments to protect cardiac tissue from the toxic effects of prolonged hypoxia or ischemia.

Cardiomyocytes (CMs), the parenchymal cell type of the heart, are particularly vulnerable to hypoxia. Despite their sensitivity to oxygen deprivation, CMs possess intrinsic adaptive mechanisms that can temporarily preserve function and viability during hypoxic stress. For instance, the fetal heart thrives in a relatively hypoxic environment. Moreover, ischemic preconditioning—brief, non-lethal episodes of ischemia—can protect the adult myocardium against subsequent severe ischemic injury.^4^ Together, these observations indicate that cardiomyocytes possess endogenous genetic programs that support survival during transient or moderate oxygen deprivation, raising the possibility that these pathways can be genetically or pharmacologically manipulated to bolster hypoxia tolerance.

Notably, cardiomyocytes are considered metabolic “omnivores,” capable of utilizing fatty acids, glucose, lactate, ketone bodies, and amino acids as energy substrates.^5^ This flexibility enables fuel switching in response to changing energetic and oxygen demands. Under normoxic conditions, fatty acid oxidation (FAO) dominates ATP production, but when oxygen is scarce, CMs metabolically rewire toward glycolysis and fuels that are less oxygen-reliant.^6^ The molecular mechanisms that enable or limit these transitions remain incompletely characterized, particularly in human CMs.

Metabolic adaptation to hypoxia has been characterized far more extensively in cancer cells and other proliferative systems. These cells optimize nutrient use to support biosynthesis and expansion of cell mass by increasing anaerobic glycolysis during hypoxia.^7^ Cardiomyocytes also shift towards fermentation when oxygen is limiting,^8^ but their metabolic priorities are fundamentally different: rather than supporting growth, they must preserve specialized functions such as electrical activity and contraction. These divergent physiological demands impose distinct constraints on how each cell type reorganizes metabolism when oxygen becomes limiting. While several genetic screens have mapped hypoxia regulators in cancer models,^9,10^ analogous efforts in cardiomyocytes have been hampered by technical challenges. Recent advances in human induced pluripotent stem cell–derived cardiomyocytes (iPSC-CMs) and CRISPR-based functional genomics now make systematic screening in CMs feasible.^11–13^

We applied these advances to systematically identify genetic regulators capable of improving cardiomyocyte tolerance to hypoxia. Using a genome-wide CRISPR screen in human iPSC-CMs exposed to varying oxygen tensions, we identified basigin (BSG) knockdown (KD) as uniquely protective in severe hypoxia. Loss of BSG led to intracellular lactate accumulation and increased pyruvate entry into the TCA cycle, thereby defending intracellular ATP levels. Our findings reveal a previously unrecognized, cardiomyocyte-specific form of metabolic regulation not observed in cancer cells or in the undifferentiated iPSC state, and suggest that modulating the transport of specific energy substrates could enhance cardiomyocyte resilience to hypoxia.

## RESULTS

### CRISPRi survival screen in iPSC-cardiomyocytes nominates basigin KD as protective in hypoxia

To identify protective targets against pathological hypoxia in cardiomyocytes, we conducted a genome-wide CRISPR inhibition (CRISPRi) screen in iPSC-CMs. We first infected iPSCs containing the CRISPRi machinery with a third-generation compact genome-wide dual-gRNA CRISPRi library^14^ (**Fig. 1a, Fig. S1a,b**) and isolated infected cells by puromycin selection (**Fig. S1c-e**). The cells were subsequently differentiated into CMs using a Wnt modulation protocol^15^ and purified via lactate purification^16^ (**Fig. S1f**). To promote maturation, we cultured the iPSC-CMs in media containing physiological levels of fatty acids, in addition to glucose, lactate, pyruvate, and other major CM carbon sources,^17,18^ given that adult CMs rely heavily on fatty acid catabolism (**Fig. S1g**). After 10 days in this “physiological” media, we confirmed increased iPSC-CM maturation by measuring elevated transcripts of key CM-specific genes relative to iPSC-CMs grown in standard high-glucose medium (**Fig. S1h**). Finally, we confirmed our population of iPSC-CMs was > 90% pure by flow cytometry using the classical CM markers, cardiac troponin T and ɑ-actinin (**Fig. S1i**).

**Figure 1.**
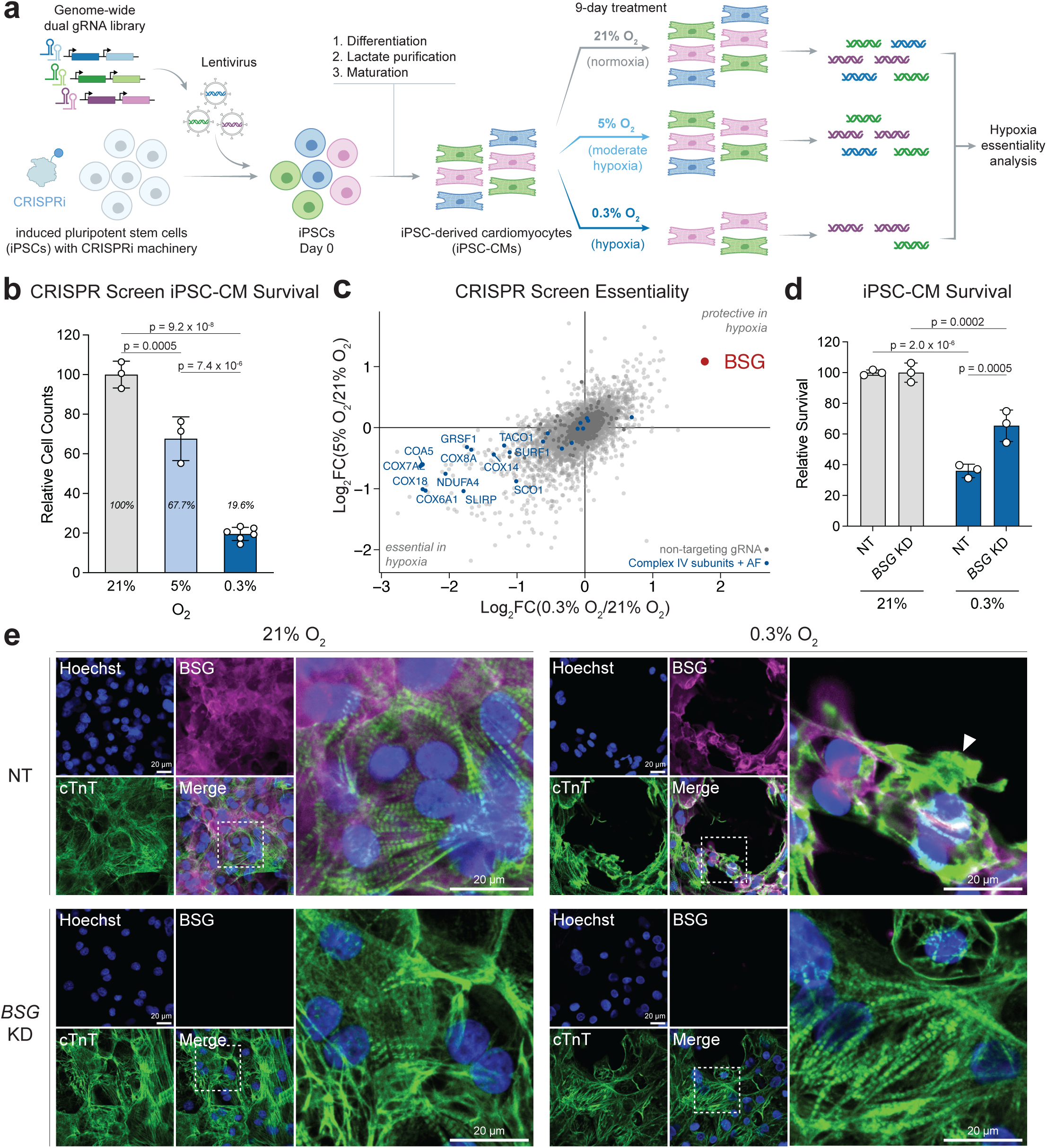
CRISPRi survival screen in iPSC-cardiomyocytes nominates basigin KD as protective in hypoxia. a, Schematic of iPSC-cardiomyocyte (CM) CRISPRi survival screen in hypoxia. After differentiation, CMs were cultured in normoxia (21% O2), moderate hypoxia (5% O2), or hypoxia (0.3% O2) for 9 days. Two independent replicates of the screen were performed. **b,** Live cell counts of iPSC-CMs from the CRISPR screen in hypoxia relative to normoxia. Each datapoint represents an individual flask. *n* = 3 replicates for 21% O2 and 5% O2, and *n* = 6 replicates for 0.3% O2. **c,** Concordance of gene essentiality between two levels of hypoxia. The protective effect of basigin (*BSG*) knockdown (KD) is highlighted as the top hit in 0.3% O₂ relative to 21% O₂. Electron transport chain (ETC) Complex IV subunits and assembly factors (AF) are indicated as essential genes across both hypoxic conditions. Non-targeting (NT) guide RNAs (gRNAs) are shown in dark grey. **d,** Validation of survival using NT or *BSG* KD iPSC-CMs cultured for 3 days in 21% O₂ or 0.3% O₂. Survival was assessed by fixing, staining, and quantifying remaining Hoechst-positive CM nuclei. *n* = 3 replicates. Values are normalized to 21% O₂ condition. **e,** Representative immunofluorescence images of NT and *BSG* KD iPSC-CMs after 3 days of culture in 21% O₂ and 0.3% O₂. Cells were stained for the nucleus (Hoechst), sarcomeres (cTnT), and BSG. The arrowhead highlights sarcomere disarray. Scale bar is 20 μm. Data are presented as mean ± SD. Significance was assessed for **b** with a one-way ANOVA with Tukey’s multiple comparisons post hoc test and for **d** with a two-way ANOVA with Tukey’s multiple comparisons post hoc test, relative to the control: NT 21% O₂.

While CMs are viable in room air (21% O2), they die over time in hypoxia.^19^ We reasoned that although cell death may not be the only or most immediate consequence of chronic hypoxia, it was the most tractable readout for a large-scale screen in post-mitotic cells such as CMs. We therefore cultured our iPSC-CMs in 21%, 5%, or 0.3% O2 for 9 days, collected the surviving CMs, and sequenced their DNA to identify gRNAs that were over- or under-represented in hypoxia relative to normoxia (**Fig. 1a**). This design allowed us to identify genes that are essential for CM survival in hypoxia (under-represented gRNAs), and also genes whose KD protects CMs from hypoxia (over-represented gRNAs). The timepoint was chosen to permit sufficient cell death for a survival-based CRISPR screen and to model chronic hypoxic stress.

As expected, more iPSC-CMs died in low oxygen, as shown by a 32% and 80% decrease in survival in 5% O2 and 0.3% O2, respectively, relative to 21% O2 (**Fig. 1b**). We performed two independent replicates of the CRISPRi screen and found a high level of concordance between the gene KDs that allowed CM survival across the two hypoxic oxygen tensions (**Fig. 1c**).

Among gene KDs with a protective effect in hypoxia, the most notable one was basigin (*BSG*) KD, the top hit in 0.3% O2 relative to 21% O2 (Log2FC ∼ 2, **Fig. 1c**). To validate this finding, we generated *BSG* KD iPSCs and differentiated them into iPSC-CMs. We observed a significant increase in survival of *BSG* KD iPSC-CMs relative to control (non-targeting (NT)) iPSC-CMs after only 3 days of culture in 0.3% O2 (**Fig. 1d**). Furthermore, *BSG* KD rescued the impaired morphology and disarrayed sarcomeres induced by extreme hypoxia (0.3% O2) (**Fig. 1e**).

Among the genes essential for survival in hypoxia, we were surprised to find genes encoding ETC Complex IV (CIV) subunits and CIV assembly factors *(GO Analysis; p = 2.2 x 10^-8^)* (**Fig. 1c, Fig. S2a**). KDs in most other ETC genes were depleted during CM differentiation or lactate purification. It is unclear why CMs with CIV KDs survived these steps. However, the dependence of CMs on CIV in hypoxia suggests that even when oxygen is limiting, iPSC-CMs are still reliant on oxidative phosphorylation.

Interestingly, existing genome-wide CRISPR screens in cancer cells did not identify basigin loss to be protective under hypoxia^9,10^ (**Fig. S2b-d**). Moreover, these screens found that ETC subunits and assembly factors are dispensable for cancer cell growth and survival in hypoxia (**Fig. S2e-g**). These observations underscore the dramatic difference in metabolic demands and vulnerabilities between differentiated cardiomyocytes and cancer cells. These differences may reflect their post-mitotic versus proliferative status, as well as intrinsic differences in metabolic wiring: cancer cells often preserve HIF-activated glycolytic pathways under normoxia,^20^ whereas cardiomyocytes depend primarily on oxidative metabolism and are particularly susceptible to hypoxic injury.

In summary, our genome-wide query of the hypoxic adaptive response identified ETC function as essential in hypoxia, and nominated *BSG* KD as protective against hypoxia-induced death in iPSC-CMs.

### *BSG* KD increases CM survival in hypoxia via loss of MCT1 and MCT4

Basigin is a highly glycosylated transmembrane protein that is primarily found in the plasma membrane.^21^ It contains two immunoglobulin-like (Ig) domains that mediate interactions with other cell-surface proteins and extracellular ligands.^22^ Basigin is widely expressed across tissues, and acts as a chaperone at the cell surface, forming complexes with cyclophilin A,^23^ matrix metalloproteinases (MMPs),^24^ as well as with monocarboxylate transporters (MCT1 and MCT4).^25^ Through these interactions, BSG regulates processes such as inflammatory signaling, extracellular matrix remodeling, and nutrient flux.

To decipher the most relevant function of BSG in hypoxic CMs, we employed a series of pharmacological inhibitors targeting these different basigin-protein interactions. Dose escalation with inhibitors of cyclophilin A (SP-8356), MMPs (AC-73), MCT1 (AZD-3965), or MCT4 (VB124) individually showed no protective effect in hypoxia. In contrast, simultaneous inhibition of MCT1 and MCT4 with syrosingopine significantly protected CMs in hypoxia in a dose-dependent manner (**Fig. 2a**). Importantly, syrosingopine is about 60-fold more potent against MCT4 (IC50 ≈ 40 nM) than MCT1 (IC50 ≈ 2.5 uM),^26^ and we only observed a rescue of CMs when the drug dose was sufficient to block both. This result suggests that pharmacological inhibition of both MCTs is necessary to phenocopy the survival benefit conferred by *BSG* transcriptional silencing. In accordance with this finding, neither *SLC16A1* (MCT1) nor *SLC16A3* (MCT4) knockdown were strongly protective hits in our CRISPR screen (**Fig. S3a**) and *SLC16A3* KD resulted in an incomplete rescue during validation experiments (**Fig. 2b-c**). Thus, the protective effects conferred by loss of basigin are likely due to disruption of both MCT1 and MCT4.

**Figure 2.**
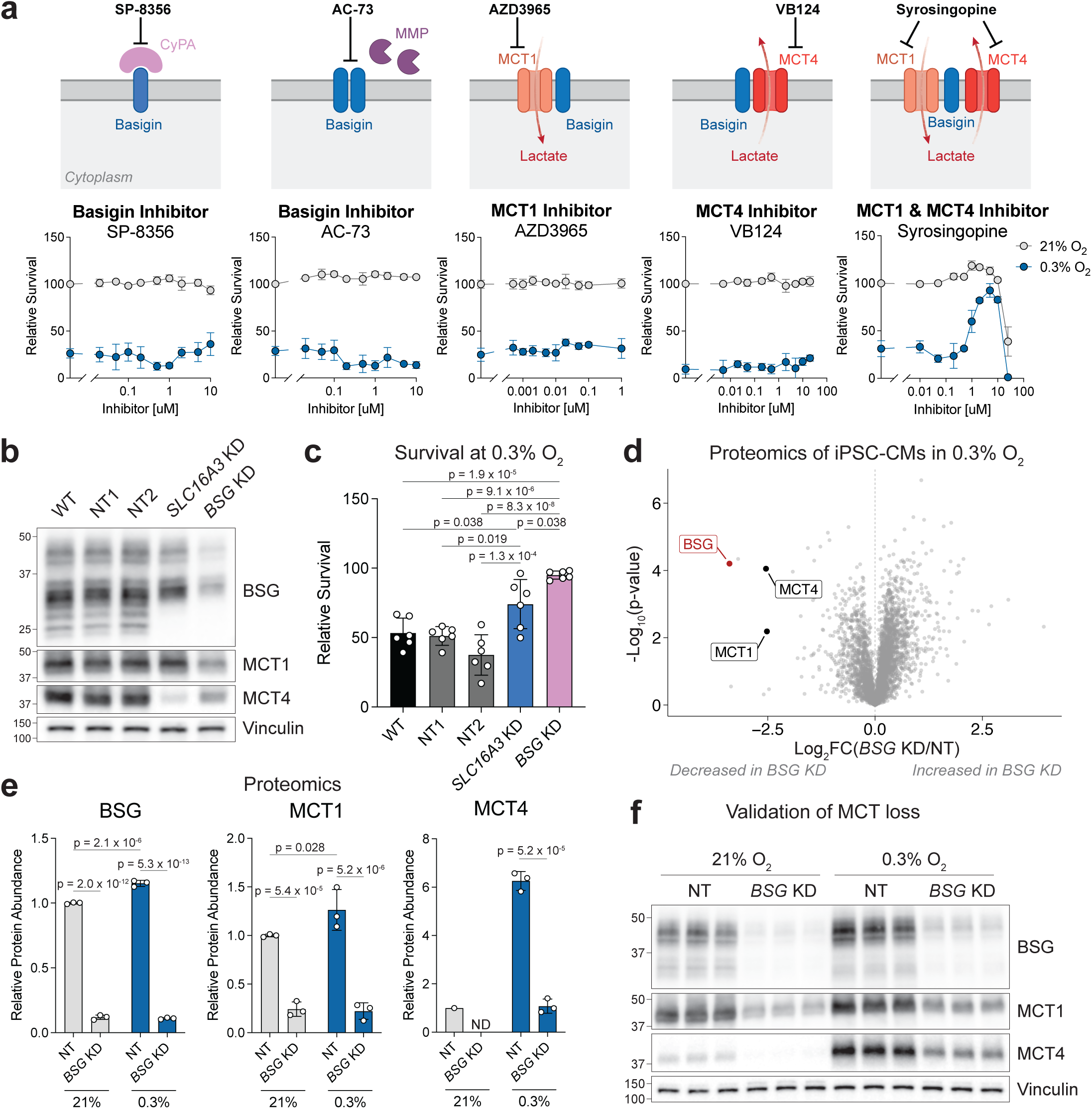
*BSG* KD increases CM survival in hypoxia via loss of MCT1 and MCT4. **a**, *Top:* Schematic of reported functions of basigin. *Bottom:* Dose-sensitivity of WT iPSC-CMs to basigin inhibitors measured after 3 days in either 21% O₂ (grey dots) or 0.3% O₂ (blue dots). *n* = 6 replicates. **b**, Western blot analysis showing levels of BSG, MCT1, and MCT4 in iPSC-CM lines, including wild-type (WT), two non-targeting controls (NT1 and NT2), *SLC16A3* (MCT4) KD, and *BSG* KD. **c**, Survival of individual KD iPSC-CMs cultured in 21% O₂ and 0.3% O₂ for 3 days. Survival data are normalized to controls in 21% O₂. *n* = 6 replicates. **d**, Proteomics of NT and *BSG* KD iPSC-CMs cultured in 0.3% O₂. *n* = 3 replicates. **e**, Proteomics quantification of BSG, MCT1, and MCT4 protein levels. ND is not detected. Statistics for MCT4 was determined with an unpaired two-tailed Student’s t-test, relative to the control: NT 0.3% O2, since MCT4 proteins were mostly undetected in 21% O2 in proteomics. **f**, Western blot of BSG, MCT1, and MCT4 protein levels in NT and *BSG* KD iPSC-CMs. Data are presented as mean ± SD. Significance was assessed for **c** with a one-way ANOVA with Tukey’s multiple comparisons post hoc test, relative to the control: WT 0.3% O2, for **d** with an unpaired two-tailed Student’s t-test, relative to the control: NT 0.3% O2, and for **e** with a two-way ANOVA with Tukey’s multiple comparisons post hoc test, relative to the control: NT 21% O₂.

Since BSG is known to be a chaperone, we investigated whether MCT levels were disrupted at the protein level in *BSG* KD iPSC-CMs. We performed proteomics under 21% and 0.3% O2 and found marked reduction of MCT1 and MCT4 protein in *BSG* KD CMs (**Fig. 2d-e**), which was confirmed by western blot (**Fig. 2f**). This is consistent with a reported role for basigin in other cell types.^22,27–29^ Moreover, we confirmed this regulation occurs post-transcriptionally, as MCT transcripts were not affected by *BSG* KD in hypoxia (**Fig. S3b-h**).

### *BSG* KD increases intracellular lactate and prioritizes glucose for pyruvate oxidation in hypoxia

The MCTs are gradient-driven, proton-coupled symporters that transport a range of substrates including pyruvate, lactate and ketones. To understand how disruption of MCT nutrient transporters could protect CMs from hypoxia, we directly measured intracellular metabolite levels by LC-MS (**Fig. 3a**). We observed an increase in intracellular lactate levels in hypoxia in NT CMs and a further increase in *BSG* KD cells in hypoxia (**Fig. 3b**). This finding was phenocopied in syrosingopine-treated NT CMs (**Fig. S4a**). In normoxia, however, when glucose is fully oxidized instead of being fermented, *BSG* KD caused minimal changes across the metabolome (**Fig. S4b,c**).

**Figure 3.**
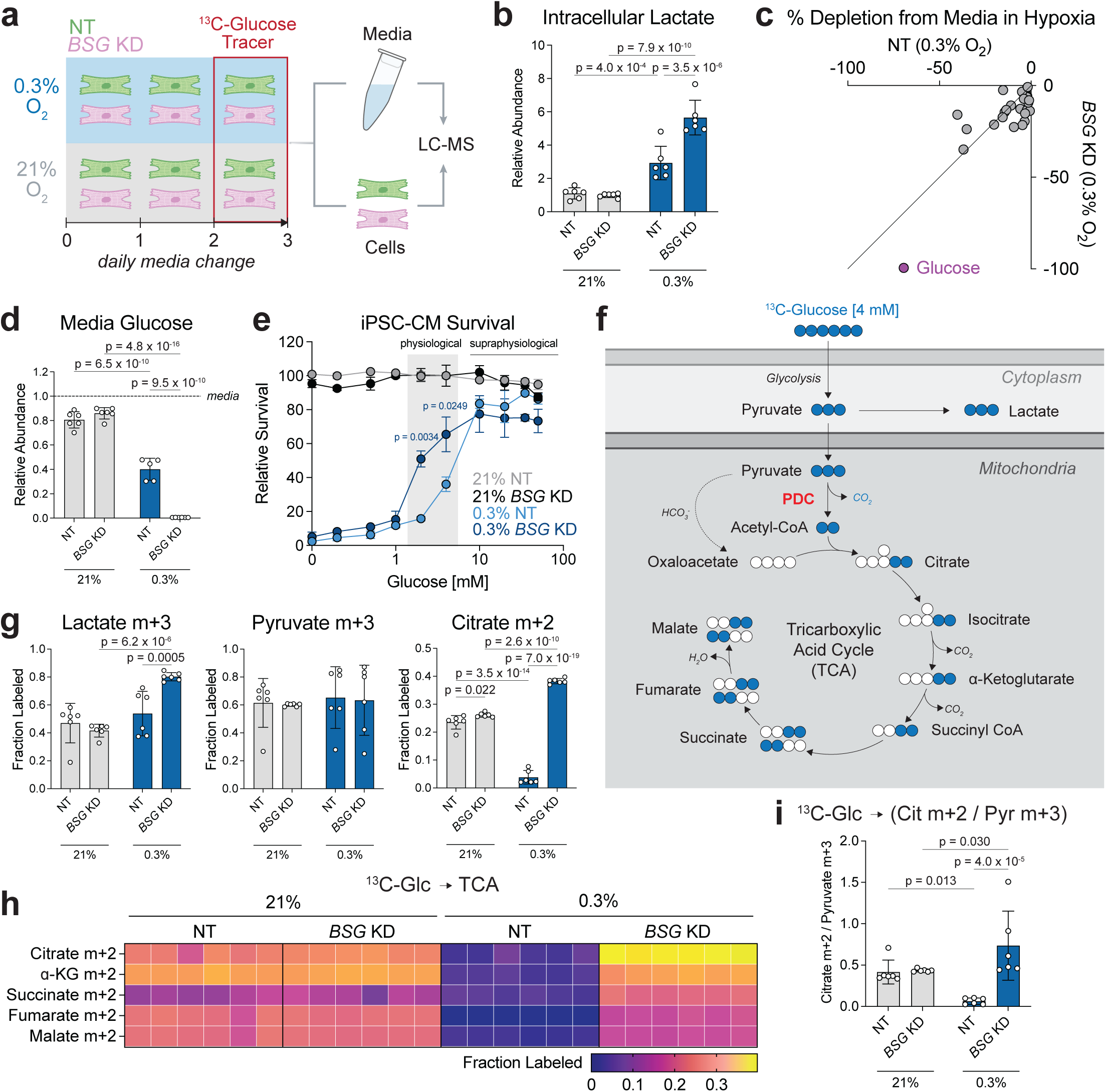
*BSG* KD increases intracellular lactate and prioritizes glucose for pyruvate oxidation in hypoxia. **a**, Schematic of metabolomics analysis of cells and conditioned media. **b**, Relative abundance of intracellular lactate (peak area normalized to an internal standard) in NT and *BSG* KD iPSC-CMs cultured in 21% O2 and 0.3% O2 for 3 days. *n* = 6 replicates. **c**, Metabolite percent depletion from media levels after 24 hr of incubation with NT and *BSG* KD iPSC-CMs at 0.3% O2. Data are means of *n* = 6 replicates. **d,** Proportion of glucose remaining in media after 24 hr of culture with NT or BSG KD CMs in 21% O2 and 0.3% O2. Normalized to the amount of glucose detected in unconditioned media. *n* = 6 replicates. **e**, iPSC-CM survival (NT vs. *BSG* KD) across a range of exogenous glucose concentrations following 3 days of culture in 21% O2 and 0.3% O2. *n* = 3 replicates. **f**, Schematic of U-^13^C-glucose tracer metabolism. U-13C-glucose carbons (blue circles) enter the TCA (via PDC, pyruvate dehydrogenase complex) or are converted to lactate. The illustrated labeling patterns show the subsequent passage of ^13^C-acetyl CoA through one round of the TCA cycle, leading to the m+2 pattern in citrate. **g**, Fraction of intracellular ^13^C-labelled to unlabeled lactate, pyruvate, and citrate after 24 hr of incubation with ^13^C-glucose media. *n* = 6 replicates. **h**, Heatmap showing the fractional labeling of all detected intracellular TCA cycle intermediates from ^13^C-glucose. *n* = 6 replicates. **i**, Ratio of intracellular citrate m+2 to pyruvate m+3 fractional labeling derived from ^13^C-glucose. *n* = 6 replicates. Data are presented as mean ± SD. Significance was assessed for **b**, **d**, **g**, and **i** with a two-way ANOVA with Tukey’s multiple comparisons post hoc test, relative to the control: NT 21% O₂.

Of note, while these transporters have biased directionality in baseline conditions (MCT1 typically imports lactate, whereas MCT4 exports it), in the context of increased lactate production, MCT1 begins to export lactate.^26,30^ In hypoxia, heightened glucose fermentation increases intracellular lactate, thus MCT1/4 loss in *BSG* KD cells leads to a net decrease in lactate export in hypoxia.

While lactate serves as a major fuel source for CMs, we were curious what other fuel sources were most consumed in hypoxia. Unsurprisingly, glucose was the most consumed metabolite in hypoxia (**Fig. 3c**). NT and *BSG* KD CMs consumed equivalent amounts of glucose in normoxia. Consistent with hypoxia increasing glycolysis,^31^ we saw an increased depletion of glucose from the media in the hypoxic vs normoxic NT cells. This effect was dramatically amplified in *BSG* KD cells, which consumed all available glucose in the media in hypoxia (**Fig. 3d**). This suggests that *BSG* KD CMs may be protected in hypoxia by increasing glucose utilization. To test if increased glucose utilization is sufficient to rescue CMs, we cultured cells for 3 days in a range of glucose concentrations and quantified CM survival. Supraphysiological levels of glucose were sufficient to rescue NT CMs (**Fig. 3e**), whereas *BSG* KD CMs survived better than NT CMs in a physiological glucose range, suggesting that glucose uptake and utilization are central to the rescue mechanism.

To determine the metabolic fate of this glucose, we tracked its relative contribution to the TCA cycle using a uniformly labeled glucose tracer (U-^13^C-glucose) (**Fig. 3f**). In the first turn of the TCA cycle, acetyl-CoA (m+2) condenses with oxaloacetate to form citrate (m+2). We observed a lower fraction of glucose-derived TCA cycle intermediates in hypoxia, again consistent with increased anaerobic glycolysis. This was completely reversed in the *BSG* KD CMs in hypoxia, where the fractional contribution of glucose to TCA cycle intermediates substantially increased (**Fig. 3g-h, Fig. S4d-f**). BSG loss therefore promotes the contribution of glucose to the TCA cycle in hypoxia.

Pyruvate is a critical branchpoint in glucose metabolism. The mitochondrial pyruvate dehydrogenase complex (PDC) is the key regulator of the oxidative versus reductive fate of pyruvate. The ratio of citrate m+2 to pyruvate m+3 provides a surrogate metric of the glucose carbons entering the TCA cycle via PDC-dependent labeling.^32,33^ Notably, *BSG* KD significantly increased this ratio, suggesting increased PDC activity in hypoxia (**Fig. 3i**). This same trend was also observed in syrosingopine-treated CMs, further indicating dual inhibition of MCT1 and MCT4 phenocopied *BSG* KD (**Fig. S4g**). Taken together, we find that *BSG* KD increases intracellular lactate, increases glucose uptake in hypoxia, and prioritizes the entry of glucose-derived carbons into the TCA cycle.

### *BSG* KD increases PDH activity by decreasing levels of PDK proteins

To explain the increased glucose contribution to the TCA cycle in *BSG* KD cells in hypoxia, we investigated the regulation of PDC activity in hypoxic CMs. Pyruvate dehydrogenase (PDH) is the central node that regulates glucose entry into the TCA cycle and is heavily regulated by different nutrient and metabolic perturbations. The E1-alpha subunit, encoded by *PDHA1*, has three primary serine (Ser) phosphorylation sites: Ser^232^, Ser^293^, and Ser^300^. Phosphorylation of any of these sites by the pyruvate dehydrogenase kinases (PDKs) suppresses PDH activity and limits glucose entry into the TCA cycle for oxidative ATP production (**Fig. 4a**). In accordance with the isotope tracer studies, we found that PDH phosphorylation of some PDHA1 sites was suppressed in *BSG* KD CMs (**Fig. 4b,c**, **Fig. S5a**), consistent with the shift towards glucose oxidation in hypoxia. To validate that the BSG-dependent decrease in PDH phosphorylation observed in iPSC-CMs extends to CMs *in vivo*, we generated a CM-specific *Bsg* deletion (*Bsg^CKO^*) in mice and found a similar decrease in p-PDH levels in the heart (**Fig. S5b,c**). This suggests that the biochemical consequences of *BSG* KD are conserved throughout diverse cardiomyocyte systems from iPSC-CMs to mouse heart.

**Figure 4.**
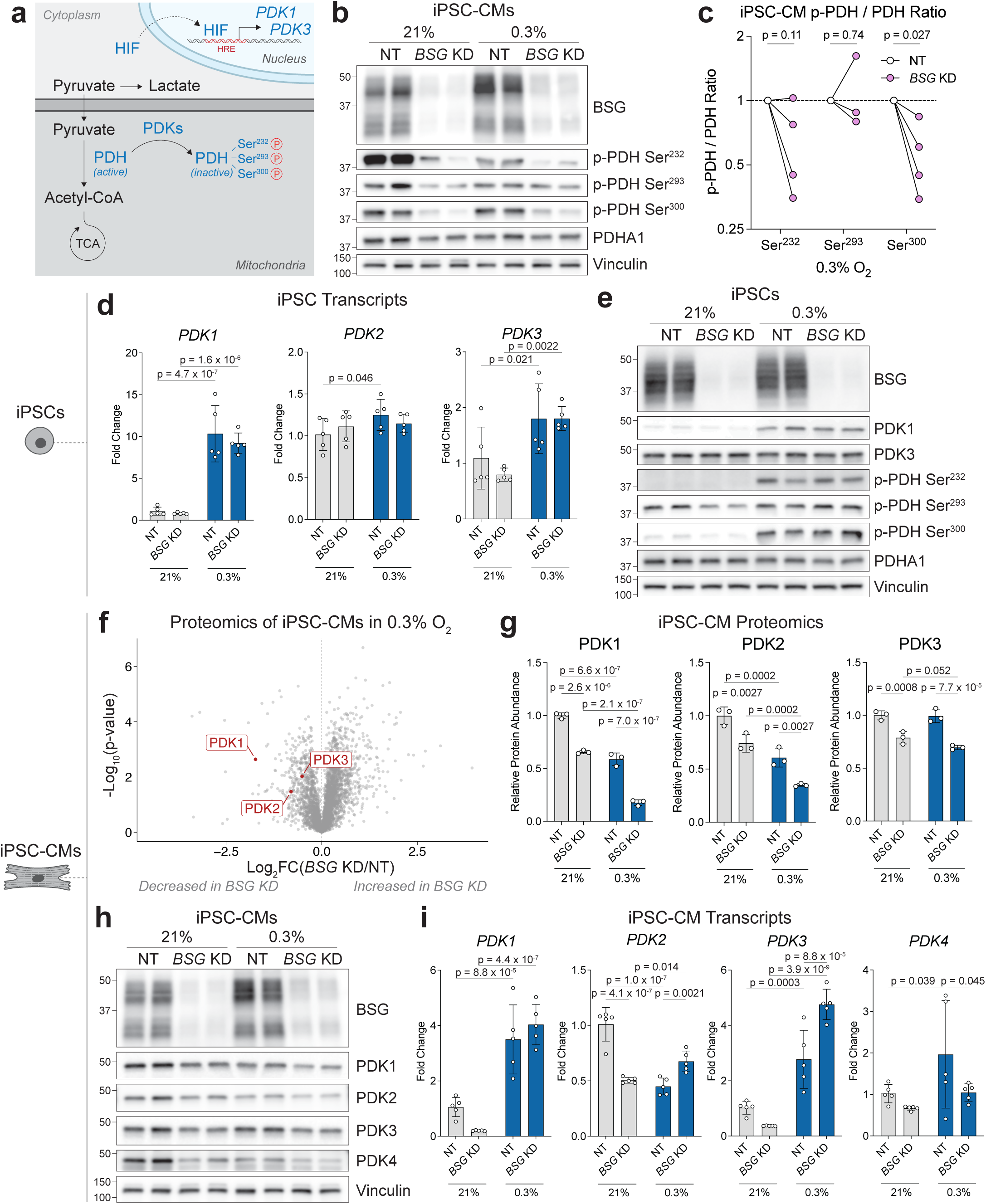
*BSG* KD increases PDH activity by decreasing levels of PDK proteins. **a**, Schematic of pyruvate dehydrogenase (PDH) regulation at the phosphorylation level by pyruvate dehydrogenase kinase (PDK). **b**, Western blot showing levels of total PDH (E1ɑ) and PDH phosphorylated at the three regulatory serine sites in NT and *BSG* KD iPSC-CMs. **c**, Phosphorylated PDH to PDH ratio in 0.3% O2 across all three serine sites, quantified from western blots from four independent iPSC-CM differentiations and hypoxia exposures. *n* = 4 replicates for Ser^232^ and Ser^300^, and *n* = 3 replicates for Ser^293^. Significance was assessed with paired t-tests comparing *BSG* KD samples with their NT controls, followed by false-discovery-rate (FDR) correction (Benjamini–Krieger–Yekutieli method). **d**, Transcriptional expression (fold change) of all detectable PDK genes measured by RT-qPCR in undifferentiated iPSCs. Values are relative to the NT 21% O2 control. *n =* 5 replicates. **e,** iPSC western blot of all detectable PDKs and phosphorylated PDH at all three serine sites. **f**, Proteomics of NT and *BSG* KD iPSC-CMs cultured in 0.3% O₂, highlighting all detected PDK proteins. Representative results of two iPSC-CM differentiations. *n =* 3 replicates. **g**, Proteomics quantification of PDK1, PDK2, and PDK3 protein levels. **h**, Western blot of PDK1, PDK2, PDK3, and PDK4 protein levels in NT and *BSG* KD iPSC-CMs cultured in 21% O2 and 0.3% O2 for 3 days. **i**, Transcriptional expression (fold change) of all PDK genes measured by RT-qPCR in iPSC-CMs. Values are relative to the NT 21% O2 control. *n =* 5 replicates. Data are presented as mean ± SD. Significance was assessed for **d**, **g**, and **i** with a two-way ANOVA with Tukey’s multiple comparisons post hoc test, relative to the control: NT 21% O2, and for **f** with an unpaired two-tailed Student’s t-test, relative to the control: NT 0.3% O2.

Importantly, hypoxia is a known regulator of PDH activity. The hypoxia responsive transcription factors (HIFs) upregulate gene expression of PDK enzymes, particularly *PDK1*.^34^ This increases inhibitory phosphorylation of PDH and decreases pyruvate oxidation when oxygen is limiting. In proliferating iPSC cells, we observed this classical adaptation where hypoxia increases *PDK1* transcript and protein levels, coupled with increased PDH phosphorylation at Ser^232^ and Ser^300^ (**Fig. 4d,e**). In iPSCs, these HIF-mediated changes in PDK and PDH phosphorylation levels were not changed in *BSG* KD cells.

Surprisingly, iPSC-CMs displayed a completely distinct pattern. By proteomics, we observed a reduction in PDK levels in hypoxia and a further reduction in *BSG* KD CMs (**Fig. 4f,g**, **Fig. S6a**). We validated this finding by western blot, showing that some PDK protein levels decreased in hypoxia and were further decreased in the *BSG* KD (**Fig. 4h, Fig. S6b**). However, the corresponding *PDK1* and *PDK3* transcripts were upregulated in CMs as they typically are under the HIF response (**Fig. 4i**). This suggests that *BSG* KD and hypoxia decrease PDKs at the protein level in CMs through post-transcriptional mechanisms. The global HIF response^35^ in CMs was unchanged by *BSG* KD (**Fig. S6c,d**), further indicating that this is a CM-specific effect on PDK protein levels, not observed in proliferating cancer cells^34^ or iPSCs.^36^ Further amplification of this effect in *BSG* KD cells was protective in hypoxia.

### Pyruvate oxidation is both sufficient and necessary for increasing CM survival in hypoxia

Our first hint that pyruvate oxidation is critical for CM survival in hypoxia came from the CRISPR screen, as ETC subunits proved particularly essential in hypoxia (**Fig. 1c**, **Fig. S2a**). To test if we could rescue NT CMs in hypoxia by forcing pyruvate oxidation, we used a pan-PDK inhibitor (dichloroacetate or DCA) to increase PDH activity (**Fig. 5a**). After 2 days of culture in 21% or 0.3% O2, a ^13^C-glucose tracer was added to NT CMs for 24 hours and the cells were maintained at their respective oxygen tensions. As expected, the total contribution of glucose to the citrate pool was lower in hypoxia and was partially reversed by DCA treatment (**Fig. 5b**). PDK inhibition was therefore sufficient to increase glucose oxidation, leading us to ask whether this alone could protect CMs. DCA has previously been shown to be protective against myocardial ischemia/reperfusion injury in rodent models,^37,38^ suggesting that it could be protective in extreme chronic hypoxia. We assessed NT iPSC-CM survival after 3 days of culture with a dose escalation of DCA in 21% and 0.3% O2. Increased PDK inhibition was <u>sufficient</u> to rescue NT CMs in hypoxia (**Fig. 5c**), indicating that forcing glucose oxidation is protective for CMs in low oxygen.

**Figure 5.**
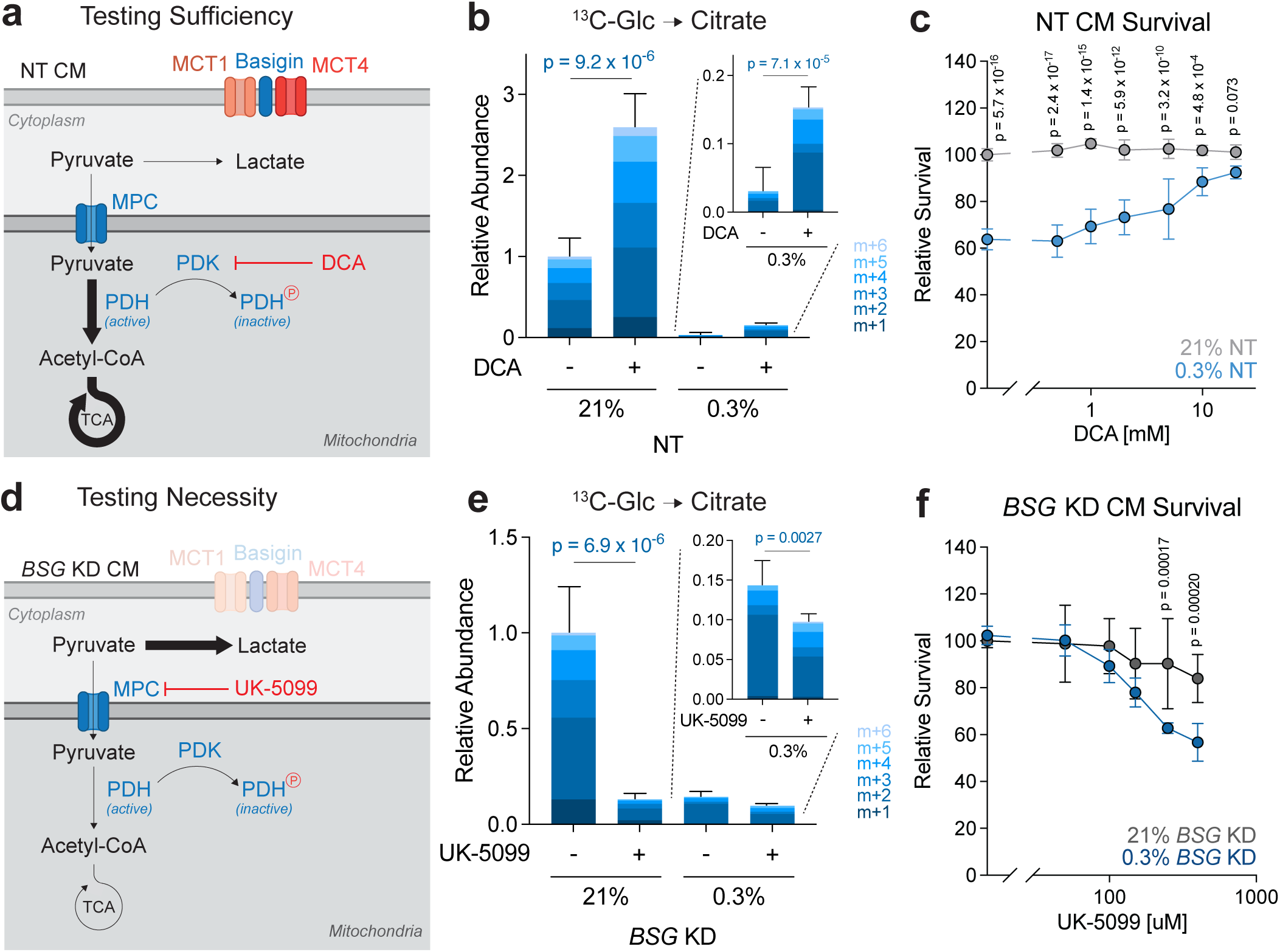
Pyruvate oxidation is both sufficient and necessary for increasing CM survival in hypoxia. **a**, Schematic of NT CMs with glucose-derived pyruvate redirected toward the TCA cycle by the addition of the PDK inhibitor DCA. **b**, ^13^C-Glucose tracer contribution to citrate ± 10 mM DCA. The graph inset shows only the samples at 0.3% O2. Statistical significance applies to differences in total labeled citrate pools. *n* = 6 replicates. **c**, Survival of NT iPSC-CMs in 0.3% O2 relative to 21% O2 as a function of PDK inhibition for 3 days with escalating DCA concentration. *n* = 6 replicates. **d**, Schematic of *BSG* KD CMs with glucose-derived pyruvate redirected toward lactate production with the addition of the mitochondrial pyruvate carrier (MPC) inhibitor UK-5099. **e**, ^13^C-glucose tracer contribution to citrate ± 250 uM UK-5099. The graph inset shows only the samples at 0.3% O2. Statistical significance applies to differences in total labeled citrate pools. *n* = 6 replicates. **f**, Survival of *BSG* KD iPSC-CMs in 0.3% O2 relative to 21% O2 as a function of MPC inhibition for 3 days with escalating UK-5099 concentration. *n* = 6 replicates. Data are presented as mean ± SD. For **b** and **e**, statistics were determined with an unpaired two-tailed Student’s t-test comparing the sums of all the labeled citrate isotopologues, and for **c** and **e**, statistics were determined with a two-way ANOVA with Šídák’s multiple comparisons post hoc test, comparing 21% O2 and 0.3 O2 at each inhibitor dose.

To test if glucose oxidation is <u>necessary</u> for the protective effect of *BSG* KD, we blocked mitochondrial pyruvate uptake by inhibiting the mitochondrial pyruvate carrier (MPC) with UK-5099 (**Fig. 5d**). As expected, blocking pyruvate uptake into the mitochondria increased whole-cell pyruvate levels (**Fig. S7b**), and as a result increased lactate production (**Fig. S7c**). With increased fermentation, we observed a concordant decrease in glucose’s contribution to the citrate pool in both normoxia and hypoxia (**Fig. 5e**). To test if blocking pyruvate oxidation affected *BSG* KD CMs, we performed a dose escalation of UK-5099 and quantified survival. *BSG* KD CMs were more sensitive to MPC inhibition in hypoxia than in normoxia, suggesting their heightened reliance on pyruvate oxidation for survival. In sum, forcing pyruvate oxidation is sufficient to protect CMs in low oxygen and is necessary for the protective effects of *BSG* knockdown in hypoxic CMs.

### Shift to pyruvate oxidation increases intracellular energy stores

We have shown that *BSG* KD CMs are protected in hypoxia by increased pyruvate oxidation, but what is the functional consequence of more pyruvate oxidation and why is it protective? One critical determinant of metabolic resilience is the NADH/NAD+ ratio. In hypoxia, the NADH/NAD+ ratio increases as NADH oxidation by the ETC decreases. As expected, we found that the NADH/NAD+ ratio increased in hypoxic NT CMs (**Fig. 6a-b**). Surprisingly, this ratio further increased in *BSG* KD CMs. The intracellular lactate/pyruvate ratio, a proxy for the cytosolic NADH/NAD+ ratio,^39^ was also higher in the *BSG* KD (**Fig. 6c**). We supplied exogenous electron donors (lactate) or acceptors (pyruvate) to modulate the NADH/NAD+ ratio, but this was insufficient to alter WT CM survival (**Fig. S8a-h**). Therefore, BSG loss does not confer protection through modulation of the NADH/NAD+ ratio, and CMs appear remarkably tolerant of hypoxia-induced shifts in both NADH/NAD+ and lactate/pyruvate redox ratios.

**Figure 6.**
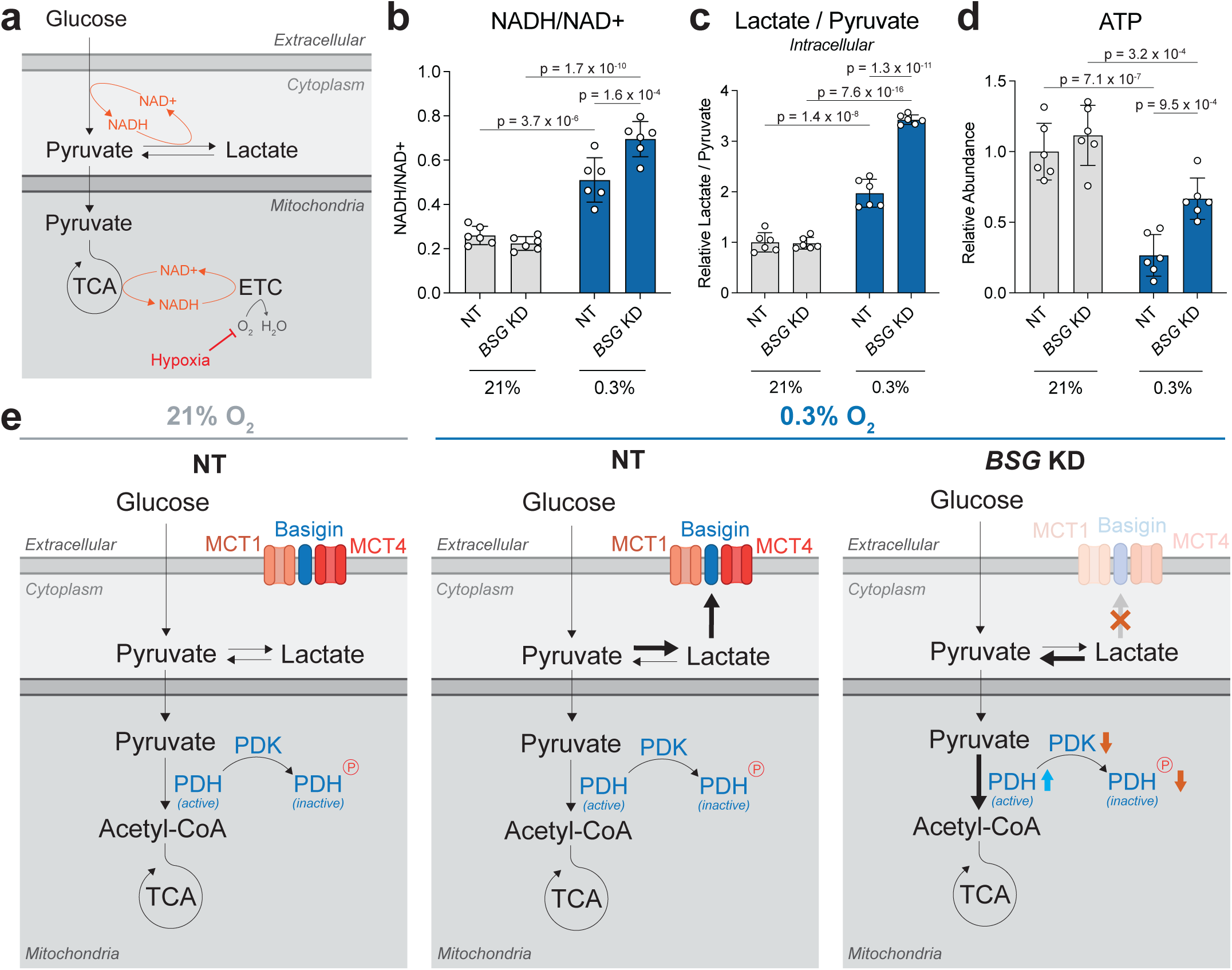
Shift to pyruvate oxidation increases intracellular energy stores. a, Diagram depicting the main NADH and NAD+ recycling reactions of central metabolism. **b,** NADH/NAD+ ratio in iPSC-CMs after 3 days in 21% O2 and 0.3% O2. *n* = 6 replicates. **c**, Intracellular lactate/pyruvate ratio, relative to NT CMs in 21% O2. *n* = 6 replicates. **d**, Intracellular ATP levels in iPSC-CMs, relative to NT CMs in 21% O2, as determined by LC-MS. *n* = 6 replicates. **e**, Schematic of the working model. Data are presented as mean ± SD. Significance was assessed for **b**-**d** with a two-way ANOVA with Tukey’s multiple comparisons post hoc test, relative to the control: NT 21% O2.

As an electrophysiological and contractile cell type, CMs have an enormous demand for ATP, so we wondered whether the loss of BSG could restore ATP levels in hypoxia. Notably, we found that hypoxic *BSG* KD CMs had significantly increased steady-state ATP levels compared to NT CMs (**Fig. 6d**). While glycolysis yields only limited ATP by converting glucose to pyruvate, complete oxidation of glucose through the TCA cycle and oxidative phosphorylation produces substantially more ATP. Pyruvate oxidation is also more oxygen-efficient than fatty acid oxidation,^40^ because β-oxidation-derived FADH₂ enters the ETC at non-proton-pumping Complex II, lowering ATP yield per O₂.^41^ Thus, when oxygen is limited, prioritizing glucose oxidation over FAO improves ATP yield per O2 molecule. Consistent with this model, we observed that total FA contribution to citrate was decreased in hypoxia using uniformly labeled U-^13^C-palmitic acid (PA) or U-^13^C-oleic acid (OA) tracers (**Fig. S9a-c**). Furthermore, *BSG* KD CMs had a lower FA incorporation into citrate compared to NT CMs, indicating preferential oxidation of glucose over FAs. Taken together, these findings demonstrate that *BSG* KD iPSC-CMs preferentially oxidize glucose in hypoxia, maintaining ATP production. In summary, our findings converge on a model in which BSG loss impairs MCT-mediated lactate export, lowers PDK abundance and PDH phosphorylation, and redirects pyruvate toward mitochondrial oxidation to maintain ATP in hypoxia (**Fig. 6e**).

## DISCUSSION

Here, we demonstrate that loss of BSG protects iPSC-CMs from pathological hypoxia. We show that *BSG* KD causes decreased MCT1/4 levels, which increases intracellular lactate. This coincides with decreased PDK protein levels, resulting in increased PDH activity. Moreover, we show that this increased glucose oxidation is both necessary and sufficient for our observed rescue effect, likely by sustaining sufficient ATP levels for cardiomyocyte survival in hypoxia. Finally, we contrast these findings to the hypoxia responses observed in iPSCs and cancer cells in which glucose oxidation is dispensable for survival in hypoxia.

Basigin is a versatile regulator of metabolism and disease across tissues. In the cardiovascular system, it contributes to pathological remodeling: heterozygous *Bsg* loss or defective BSG glycosylation reduces cardiac fibrosis and preserves ventricular function in pressure-overload and transverse aortic constriction models.^42,43^ Cyclophilin A–BSG signaling also drives hypoxia-induced pulmonary hypertension by promoting inflammation, smooth muscle proliferation, and vascular remodeling, and *Bsg* heterozygosity attenuates these responses.^44^ Consistent with these findings, a human *BSG* polymorphism associated with lower *BSG* transcript levels reduces the risk of chronic heart failure.^45^ In cancer and immune cells, BSG loss suppresses glycolytic flux and inhibits tumor growth^46^—mirroring the effects of dual MCT1/MCT4 inhibition with agents like syrosingopine.^47^ Together, these observations underscore basigin as a context-dependent regulator of nutrient handling and stress responses.

While it is counterintuitive that impaired lactate export could be beneficial for CMs in hypoxia, recent work has demonstrated that MCT disruption is protective against cardiac injury. Specifically, pharmacological inhibition of MCT4 has been shown to prevent isopreterenol-induced cardiac hypertrophy in a murine heart failure model,^48^ and has improved cardiac function after ischemia-reperfusion injury in mice.^49^ This is consistent with the partial protection we observed with an *SLC16A3* (MCT4) KD in iPSC-CMs in hypoxia. Interestingly, the loss of MCT1 has been shown to worsen heart failure in mice.^50^ In skeletal muscle however, deletion of *Slc16a1* (MCT1) in mice increases glucose contribution to the TCA and promotes formation of oxidative myofibers.^51^ In iPSC-CMs, we find that dual inhibition of MCT1 and MCT4 is protective in hypoxia. These subtle differences may reflect varying degrees and durations of hypoxia across experimental systems. Further work will be needed to dissect the contribution of each MCT to hypoxic cardiac pathology across cell states and different hypoxic and ischemia/reperfusion regimes.

The canonical HIF response upregulates *SLC16A3* (MCT4) to facilitate lactate export.^52^ While this may be acutely protective in CMs, it may become maladaptive over prolonged hypoxia. Unlike iPSCs which rely more heavily on glycolysis and therefore tend toward lactate export, CMs primarily oxidize lactate as an energy substrate. Consistent with these distinct metabolic tendencies, the transcriptional induction of *SLC16A3* (MCT4) under hypoxia was over 5-fold stronger in iPSCs than in iPSC-CMs. HIF also classically upregulates *PDK1* in hypoxia, and this response was similarly over 2-fold greater in iPSCs, suggesting that pluripotent cells mount a more pronounced transcriptional response for lactate production and export under hypoxic conditions.

In CMs, many of the PDKs are already present in normoxia, consistent with reports that at baseline, only about 61% of the PDH in the heart is in its active (de-phosphorylated) form.^53^ This low baseline is hypothesized to enable the heart to rapidly respond to stressors or increased workload to expand the capacity for pyruvate oxidation. In hypoxia, *BSG* KD CMs displayed increased pyruvate oxidation, and in line with this, we found that PDK protein levels were reduced in the KD condition. This effect was not observed in iPSCs, suggesting a CM-specific regulation of PDK protein levels. PDK stability could be affected by many factors, including some of the secondary consequences of BSG loss such as increased lactate or potentially low pH. In line with this, a group has recently shown that PDK3 enzyme structure and activity is sensitive to slightly acidic conditions,^54^ though this link warrants further investigation.

Additionally, lactate accumulation in *BSG* KD CMs may result in increased LDH activity towards pyruvate due to mass action principles, thereby increasing glucose and pyruvate entry into the TCA cycle. Furthermore, recent work has shown that lactate can directly activate the ETC independently of its metabolism,^55^ raising the possibility that elevated intracellular lactate in *BSG* KD CMs could augment ETC activity. Future work will be needed to dissect the relative contributions of these additional mechanisms to the *BSG* KD rescue effect in hypoxia.

CMs exhibit metabolic flexibility, with glucose and fatty acid oxidation (FAO) reciprocally regulated by the Randle cycle.^56^ PDKs are key regulators of this fuel selection, and it has been reported that *Pdk1* deletion in the heart not only increases glucose oxidation but decreases FAO,^57^ whereas increased *Pdk4* expression enhances FAO.^58,59^ Consistent with these reports, we observed that *BSG* KD CMs have lower fatty acid incorporation into the TCA cycle (**Fig. S8a-c**). We have recently shown that the heart shifts toward glucose oxidation and away from FAO in response to chronic hypoxia.^60,61^ During FAO, β-oxidation produces FADH₂ in addition to NADH. Because FADH₂ donates electrons to Complex II, which does not pump protons, FAO yields fewer ATP molecules per O₂ consumed compared to glucose oxidation.^40,41^ Consequently, switching to glucose oxidation is considered O₂-sparing and protective under hypoxic conditions. As we and others have described, pharmacologically forcing pyruvate oxidation during chronic hypoxia or ischemia-reperfusion with DCA is protective.^37,38^ *BSG* KD could potentially facilitate a similar metabolic shift by promoting pyruvate entry into the TCA cycle.

How is forcing pyruvate oxidation helpful even when oxygen is limited? Cytochrome c oxidase (Complex IV) of the ETC has a very high affinity for oxygen, with a Km of 0.1% O2.^62,63^ Therefore, in states of hypoxia (not anoxia), Complex IV still has the ability to bind oxygen. In hypoxia, the HIF response may be prematurely shutting down oxidative ATP production and contributing to CM death. In our CRISPRi screen, we identified Complex IV as essential in low oxygen, reinforcing that a functional ETC is still critical at this oxygen-tension. In this vein, MPC loss has been shown to promote cardiac hypertrophy^48^ and to exacerbate infarct size in a myocardial infarction model.^49^ Pyruvate oxidation is therefore paramount to generate sufficient energy for the heart.

Finally, our finding that steady-state ATP levels were higher in *BSG* KD CMs in hypoxia even when the NADH/NAD+ ratio was high, implies that maintaining energetics is more important than redox balance in CMs. This is in contrast to proliferating cells like iPSCs and cancer cells in which the demand to regenerate NAD+ can sometimes surpass the demand for ATP.^64^ In cancer cells, even the ETC can become dispensable for proliferation, as long as NAD+ regeneration and nucleotide synthesis can occur.^65^ Meanwhile, CMs are unable to sustain viability on anaerobic glycolysis, even when excess pyruvate is provided to WT CMs as an electron acceptor to regenerate NAD+.

More broadly, our findings identify basigin as a novel protective target in cardiac hypoxia and nominate dual MCT1/MCT4 inhibition as a promising strategy for future *in vivo* studies. These results also highlight the need for more effective approaches to enhance glucose oxidation. Although dichloroacetate (DCA) has shown efficacy in rodent models,^37,38^ its clinical translation has been limited by dose-dependent, reversible peripheral neuropathy.^66^ Thus, the development and testing of safer, more potent activators of this pathway—ideally in human-based cardiac models—will be critical. Targeting MCTs and/or basigin may offer an alternative means to promote pyruvate oxidation in cardiomyocytes.

## ACKNOWLEDGEMENTS/FUNDING

We thank all the members of the Jain Lab for their thoughtful discussion and review of the manuscript. We thank the Nakamura Lab for providing WTC-11 iPSCs with the dCas9-KRAB machinery. We thank Dr. Joseph Replogle for providing CRISPRi plasmids and help with library preparation. We thank Dr. Gabriela Grigorean at UC-Davis for assistance with proteomics. We give our deepest thanks to the Gladstone Cores: Stem Cell Core, Genomics Core, Flow Cytometry Core, Assay Development and Drug Discovery Core, and Histology and Light Microscopy Core. We thank Dr. Françoise Chanut and Dr. Brian Plosky for their insights during the writing process and Dr. Chiara Ricci-Tam for help with figure generation. Sequencing was performed at the UCSF CAT, supported by UCSF PBBR, RRP IMIA, and NIH 1S10OD028511-01 grants. Funding was provided by the National Heart, Lung, and Blood Institute F31HL172581 (W.R.F.), National Institute on Aging R01AG065428 (W.R.F., I.H.J, and K.N.), the National Institute of General Medical Sciences Medical Scientist Training Program, grant T32GM141323 (A.D.M.), the National Institute of Diabetes, Digestive, and Kidney Diseases under grant 5F30DK139713 (A.D.M.), the National Heart, Lung, and Blood Institute K08HL169915 (A.H.B.), the American Heart Association Harold Amos Medical Faculty Development Award (A.H.B.), Keck Medical Research Grant (I.H.J.) and Arc Institute support (I.H.J. and G.S.).

Any opinions, findings, and conclusions or recommendations expressed in this material are those of the authors and do not necessarily reflect the views of the National Institutes of Health.

## AUTHOR CONTRIBUTIONS

W.R.F. and I.H.J. conceived the project. W.R.F. designed and conducted experiments and analyzed the data. A.D.M. and S.Y.M. assisted with stable isotope tracer study design and analysis. A.D.M., S.Y.M., Y.M.M., M.W.C., Y.H., A.H.B., H.H., G.S., and N.K.B. assisted with sample collection. R.A.N. provided reagents/materials. D.S. and K.N. helped supervise the research. W.R.F. and I.H.J. wrote the manuscript, with input from all authors.

## CORRESPONDING AUTHORS

Correspondence to Isha H. Jain and Ken Nakamura

## ETHICS DECLARATIONS

I.H.J. has patents related to hypoxia therapy.

## CODE/DATA AVAILABILITY

Cell lines and plasmids can be made available from the corresponding author upon request and MTA.

## SUPPLEMENTARY FIGURES

**Figure S1.**
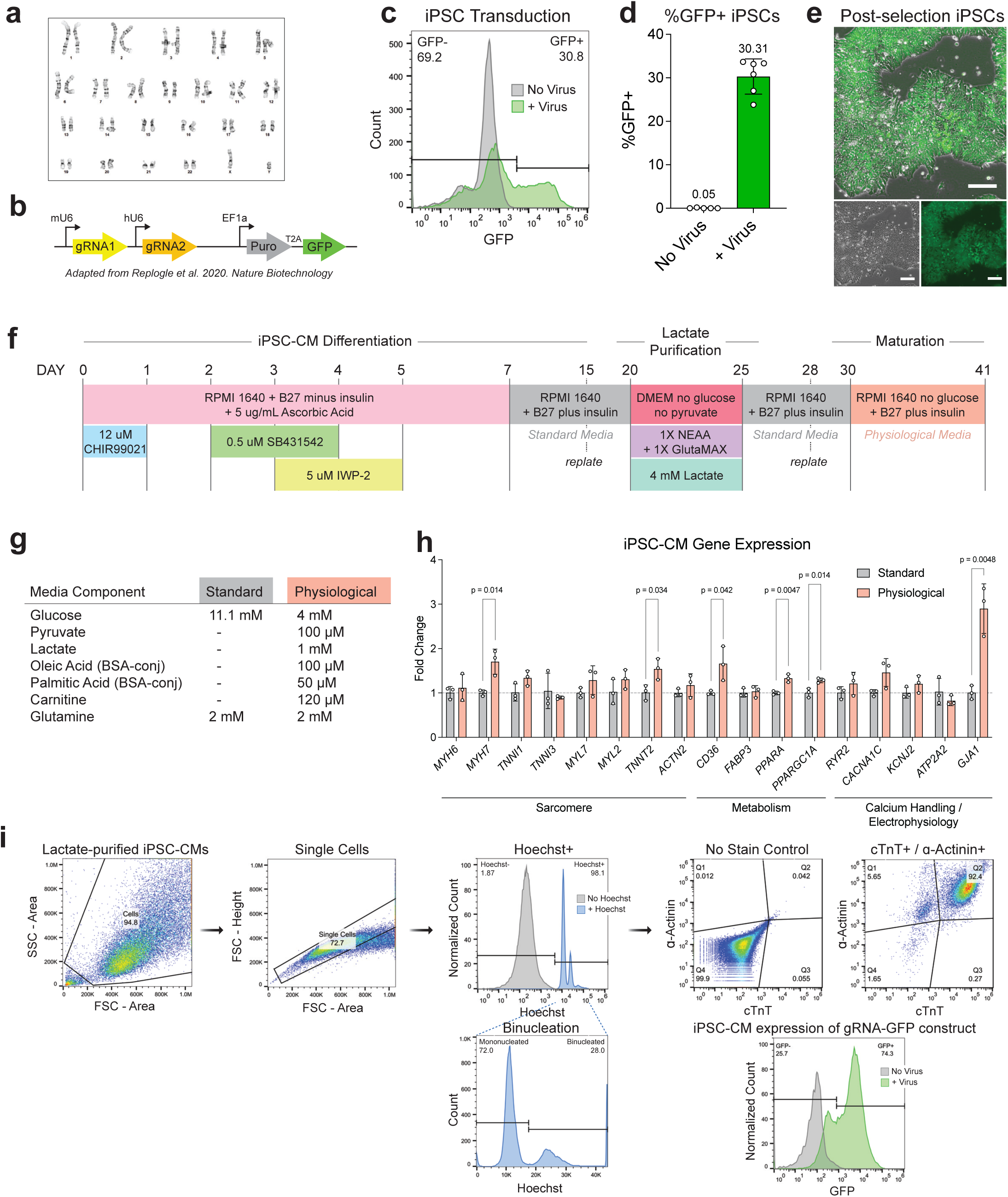
Genome-wide CRISPRi survival screen in chronic hypoxia in iPSC-CMs methods. **a**, Normal karyotype of WTC-11 iPSCs with dCas9-KRAB in the CLYBL locus. **b**, Diagram of the dual gRNA cassette from the genome-wide CRISPRi library with each gRNA driven by the mouse (mU6) or human (hU6) U6 promoter. Each construct contains puromycin resistance and GFP driven by EF1α. **c**, Representative lentiviral infection of genome-wide gRNA library into dCas9-KRAB iPSCs at a target MOI of 0.3. GFP measurement via flow cytometry enabled quantification of the percent of iPSCs infected. **d**, MOI infection of each T75 flask of iPSCs from the CRISPRi screen. *n* = 5 replicates. **e**, GFP+ iPSCs with genome-wide gRNA library after puromycin selection. Scale bar is 100 μm. **f**, Diagram of iPSC-CM differentiation protocol. Differentiation was initiated with Wnt signaling induction using the GSK3 inhibitor CHIR99021, followed by TGF-β/activin/NODAL pathway inhibition with the ALK4/5/7 inhibitor SB431542, and finally Wnt pathway inhibition with Porcupine inhibitor IWP-2. RPMI 1640 with B27 without insulin was supplemented with Ascorbic Acid to increase iPSC-CM differentiation efficiency. RPMI 1640 with B27 plus insulin (standard media) was changed on day 7, and iPSC-CMs were re-plated on day 15 for lactate purification. iPSC-CMs were replated again on day 28 after purification and were matured in physiological media for 10 days. **g**, Media compositions of standard high glucose media and physiological media used in this study. **h**, Transcriptional expression (fold change) of cardiac maturation marker genes measured by RT-qPCR in iPSC-CMs cultured in either standard or physiological media for 10 days. *n* = 3 replicates. Statistical significance was evaluated using a two-tailed, unpaired Student’s t-test, relative to standard media. **i**, Lactate purified iPSC-CMs containing the CRISPRi library were analyzed by flow cytometry. Binucleation was detectable in 28% of iPSC-CMs, and 92.4% of cells were double-positive for cTnT and α-Actinin.

**Figure S2.**
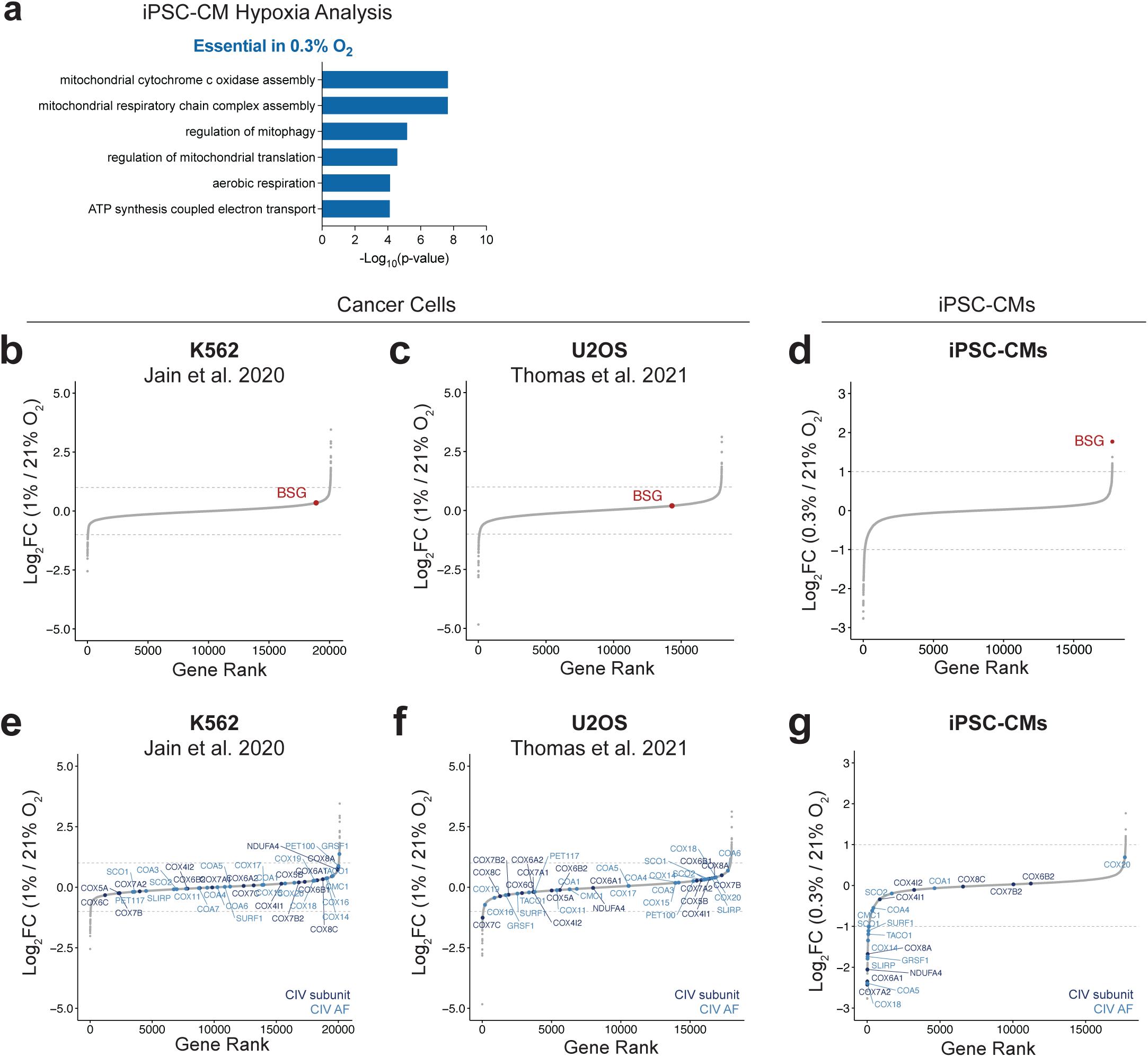
CRISPRi screen in iPSC-CMs identifies cardiomyocyte-specific responses to hypoxia. **a**, GO terms from hits from iPSC-CM CRISPRi screen of 0.3% vs 21% O₂. The top GO term shows Complex IV essentiality in hypoxia: “mitochondrial cytochrome c oxidase assembly” (GO:0033617). Other top GO terms show ETC essentiality in hypoxia, including: “mitochondrial respiratory chain complex assembly” (GO:0033108), “aerobic respiration” (GO:0009060), and “ATP synthesis coupled electron transport” (GO:0042773). **b, c**, Genome-wide CRISPRn screens in K562 (**b**) and U2OS (**c**) cancer cells did not find *BSG* deletion to be protective in hypoxia (1% O₂) relative to normoxia (21% O₂). **d**, CRISPRi screen in iPSC-CMs showing increased survival of *BSG* KD CMs in hypoxia (0.3% O₂) relative to normoxia (21% O₂). **e, f**, Genome-wide CRISPRn screens in K562 (**e**) and U2OS (**f**) cancer cells do not find Complex IV subunits or assembly factors to be essential in hypoxia (1% O₂) relative to normoxia (21% O₂). **g**, CRISPRi screen in iPSC-CMs showing increased essentiality of Complex IV subunits and assembly factors in hypoxia (0.3% O₂) relative to normoxia (21% O₂).

**Figure S3.**
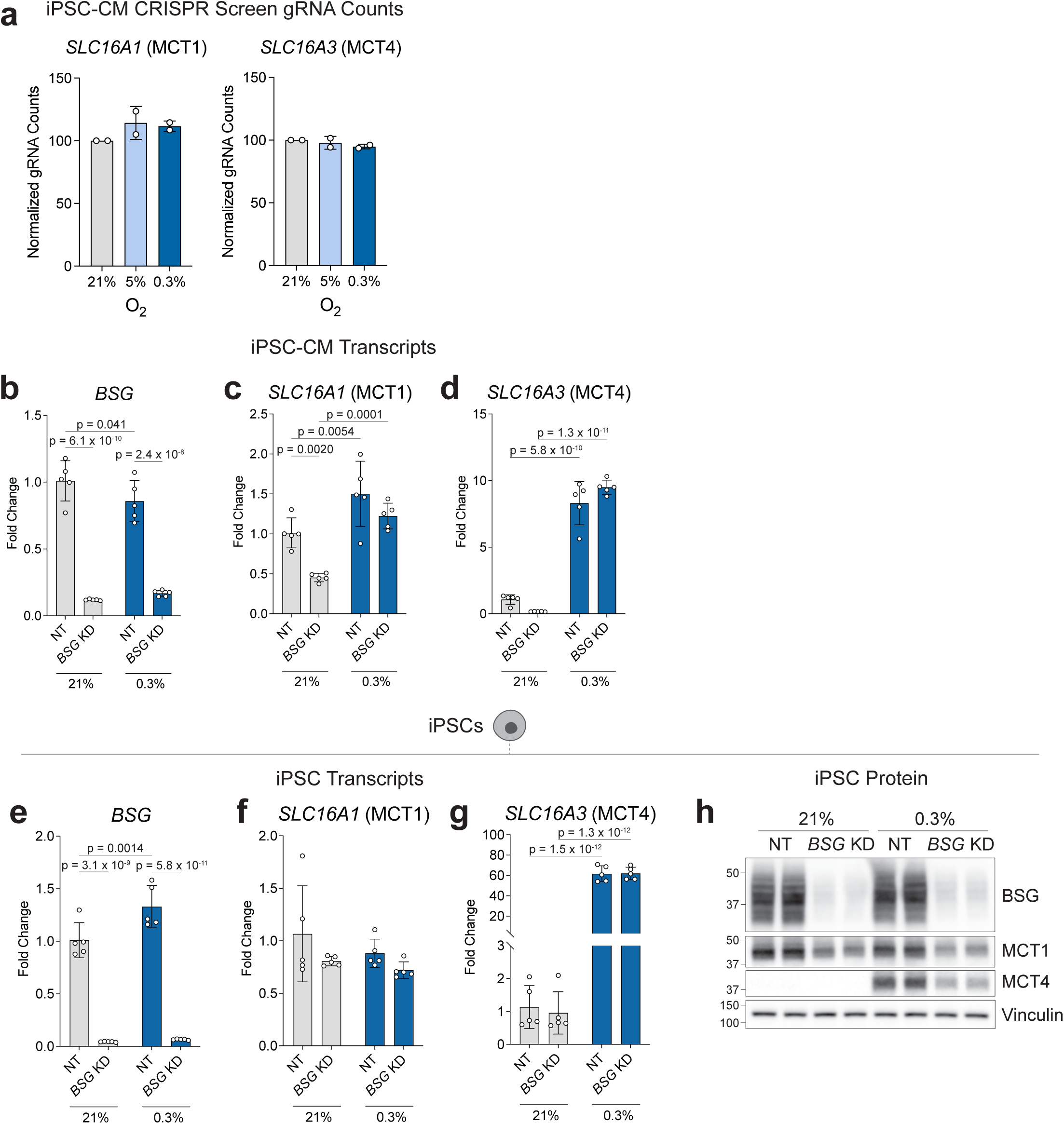
MCT1 and MCT4 are regulated by BSG at the protein level in hypoxia. **a**, gRNA counts from the two replicates of the CRISPRi screen for *SLC16A1* (MCT1) KD and *SLC16A3* (MCT4) KD, normalized to gRNA counts at 21% O2. **b**, Fold change of iPSC-CM transcripts, relative to NT at 21% O2 after 3 days of culture. Transcripts measured by RT-qPCR for *BSG* (**b**), *SLC16A1* (**c**), and *SLC16A3* (**d**). *n* = 5 replicates. **e**, Fold change of iPSC transcripts, relative to NT at 21% O2 after 3 days of culture. Transcripts measured by RT-qPCR for *BSG* (**e**), *SLC16A1* (**f**), and *SLC16A3* (**g**). *n* = 5 replicates. **h**, Western blot of NT and BSG KD iPSCs after 3 days of culture at 21% O2 and 0.3% O2, blotting for BSG, MCT1, and MCT4 levels. Data are presented as mean ± SD. Significance was assessed for **b**-**g** with a two-way ANOVA with Tukey’s multiple comparisons post hoc test, relative to the control: NT 21% O2.

**Figure S4.**
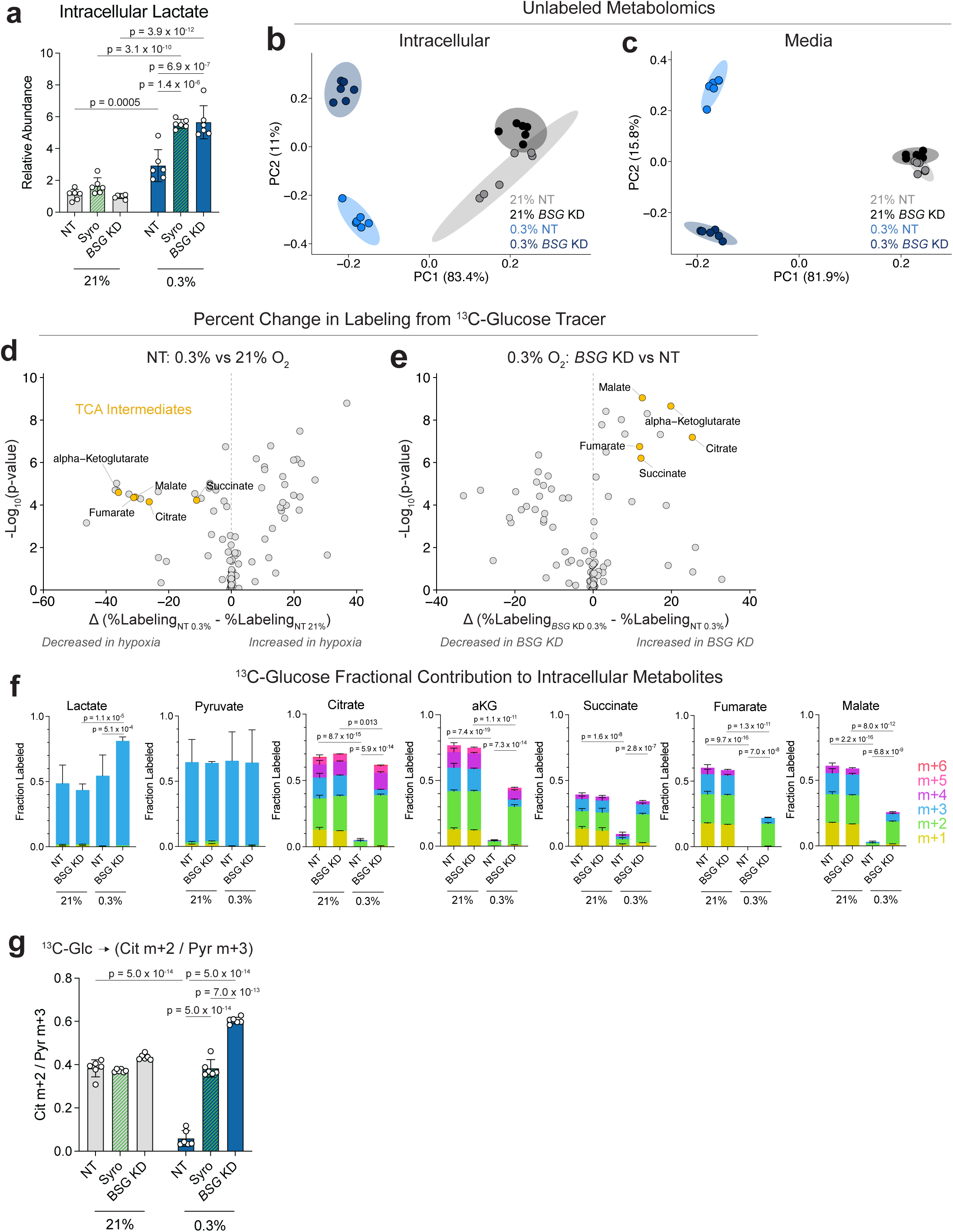
*BSG* KD increases glucose contribution to TCA in hypoxia. **a**, Relative abundance of intracellular lactate in iPSC-CMs cultured 21% O2 and 0.3% O2 for 3 days. Treatment conditions include NT, NT plus 2 μM syrosingopine, and *BSG* KD CMs, with data normalized to the NT 21 O2 condition. *n* = 6 replicates. **b**, PCA analysis of all intracellular metabolites across NT and *BSG* KD CMs after 3 days of culture in 21% O2 and 0.3% O2. Ellipses represent 95% confidence intervals within each group. The axes show the percent of variation explained by each principle component (PC). Analysis is representative of 145 metabolites detected by LC-MS. *n* = 6 replicates. **c**, PCA analysis of all metabolites detected in 24-hour conditioned media from NT and *BSG* KD CMs after 3 days of culture in 21% O2 and 0.3% O2. Analysis is representative of 130 metabolites detected by LC-MS. *n* = 6 replicates. **d**, ^13^C-glucose tracer contribution to all detected metabolites, shown as the change in percent labeled carbons of each metabolite between hypoxic (0.3% O2) and normoxic (21% O2) conditions. TCA cycle intermediates are highlighted. **e**, ^13^C-glucose tracer contribution to all detected metabolites, shown as the change in percent labeled carbons of each metabolite between *BSG* KD CMs in 0.3% O2 and NT CMs in 0.3% O2. **f**, ^13^C-glucose tracer fractional contribution to lactate, pyruvate, and TCA intermediates. Statistics compare the sum of glucose-derived isotopomers across conditions. *n* = 6 replicates. **g**, ^13^C-glucose tracer contribution to citrate m+2 / pyruvate m+3 after 3 days of culture in 21% O2 or 0.3% O2. 2 μM syrosingopine was included for 3 days on NT CMs as a comparison to *BSG* KD CMs. *n* = 6 replicates. Data are presented as mean ± SD. Significance was assessed for **a** and **g** with a one-way ANOVA with Tukey’s multiple comparisons post hoc test, relative to the control: NT 21% O2, and for **f** with a two-way ANOVA with Tukey’s multiple comparisons post hoc test, relative to the control: NT 21% O2.

**Figure S5.**
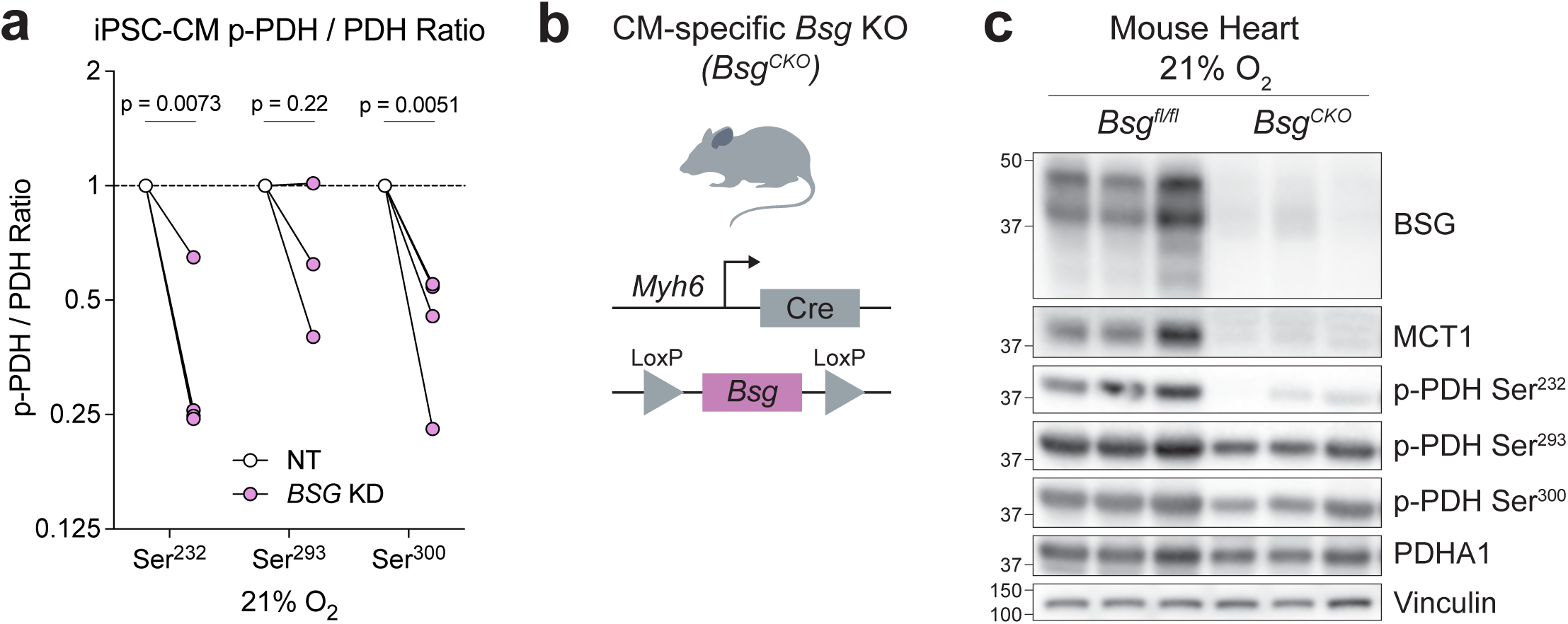
BSG deficiency reduces PDH phosphorylation in human iPSC-CMs and adult murine CMs. **a**, Ratio of phosphorylated PDH (p-PDH) to total PDH across three inhibitory serine sites in iPSC-CMs under normoxia (21% O2). Values were quantified via western blot from four independent iPSC-CM differentiations. *n* = 4 replicates for Ser^232^ and Ser^300^, and *n* = 3 replicates for Ser^293^. Significance was assessed with paired t-tests comparing *BSG* KD samples with their NT controls, followed by false-discovery-rate (FDR) correction (Benjamini–Krieger–Yekutieli method). **b**, Schematic for the generation of a cardiomyocyte-specific conditional *Bsg* knockout (*Bsg^CKO^*) mouse model, using a *Bsg^fl/fl^*;*Myh6*-Cre driver. **c**, Western blot analysis of BSG, MCT1, PDHA1, and p-PDH (Ser^232^, Ser^293^, Ser^300^) protein levels in heart tissue from WT (*Bsg^fl/fl^*;*Myh6*-Cre^-^) and *Bsg^CKO^* (*Bsg^fl/fl^*;*Myh6*-Cre^+^) mice. *n* = 3 mice per group.

**Figure S6.**
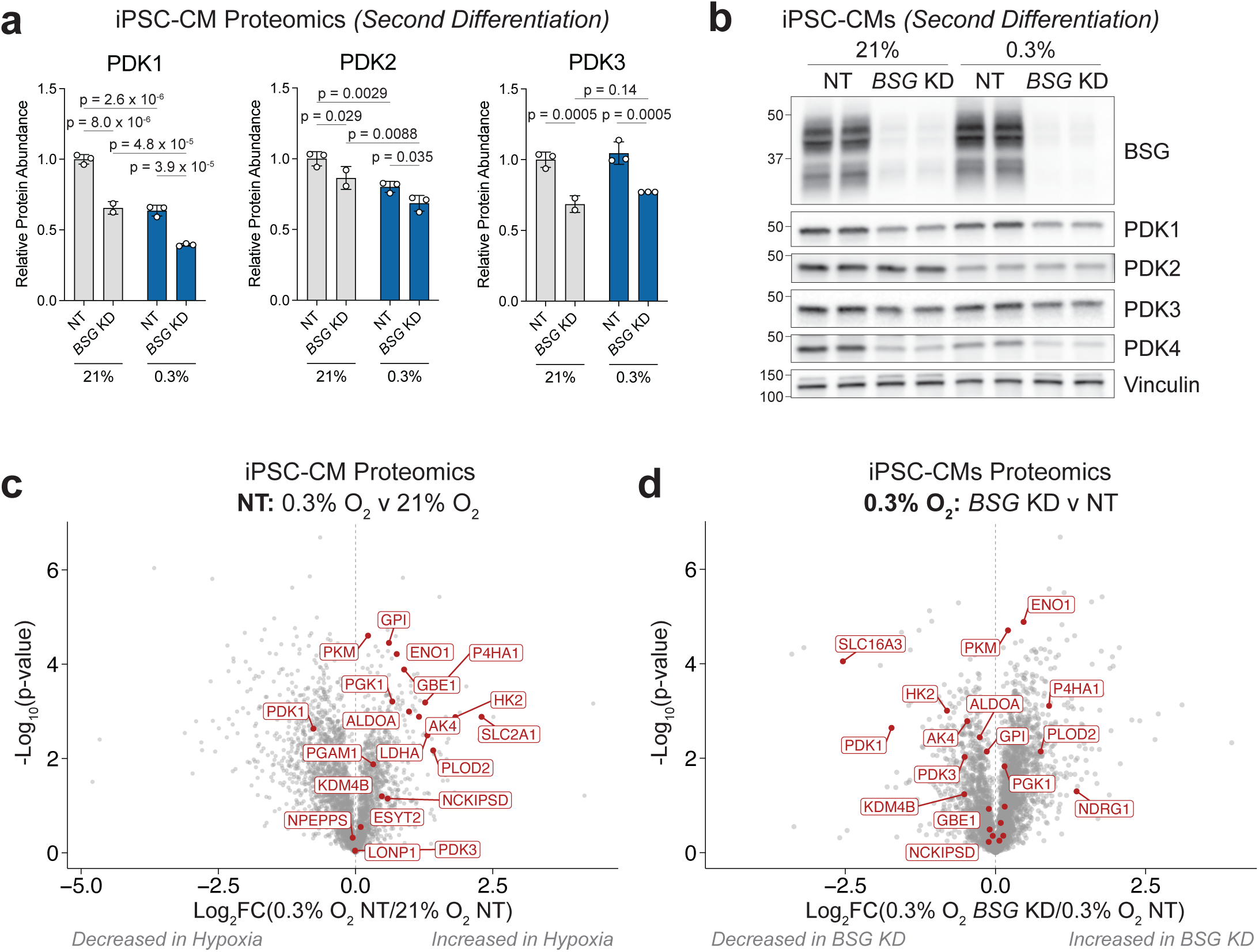
*BSG* KD does not affect global HIF target protein abundance. **a**, Proteomics of NT and *BSG* KD CMs from a second iPSC-CM differentiation, highlighting all detected PDK proteins. *n =* 3 replicates. **b**, Western blot of PDK1, PDK2, PDK3, and PDK4 protein levels in NT and *BSG* KD iPSC-CMs cultured in 21% O2 and 0.3% O2 for 3 days. From an independent iPSC-CM differentiation and hypoxia exposure. **c**, Proteomics of NT CMs in 0.3% O2 versus 21% O2, highlighting known HIF targets. *n* = 3 replicates. **d**, Proteomics of NT versus *BSG* KD CMs in 0.3% O2, highlighting known HIF targets. *n* = 3 replicates. Data are presented as mean ± SD. Significance was assessed for **a** with a two-way ANOVA with Tukey’s multiple comparisons post hoc test, relative to the control: NT 21% O2, and for **c** and **d** with an unpaired two-tailed Student’s t-test, relative to the controls: NT 21% O2 and NT 0.3% O2, respectively.

**Figure S7.**
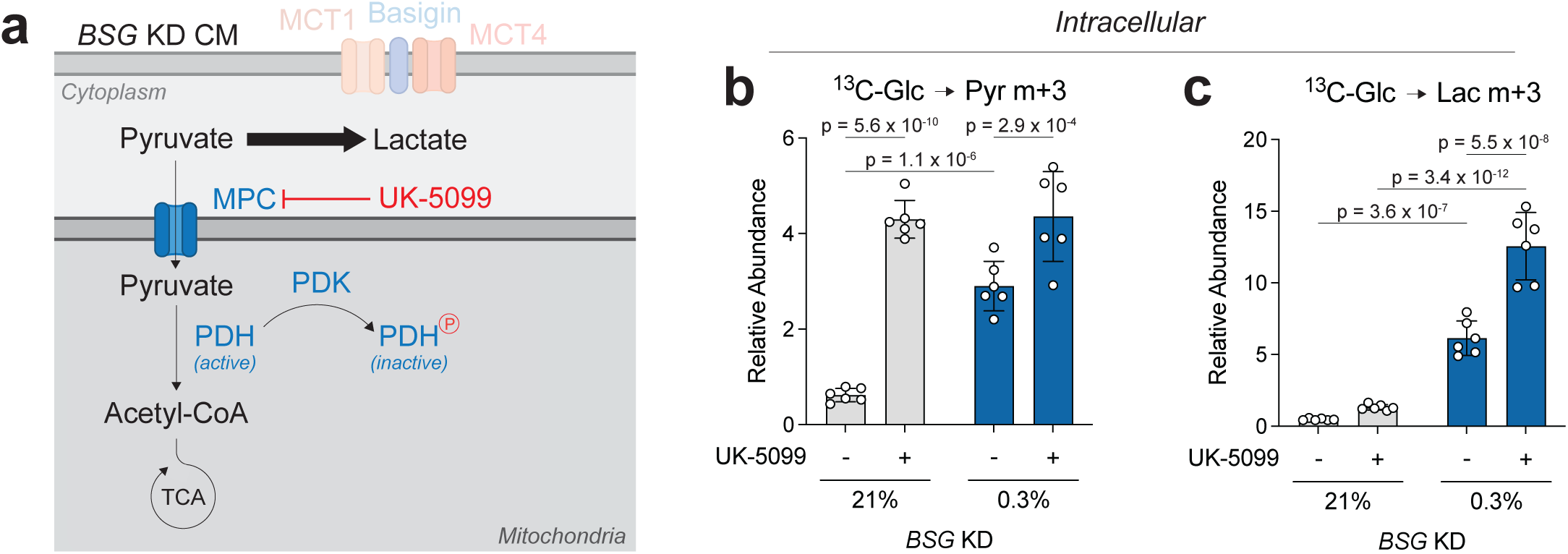
Inhibiting mitochondrial pyruvate uptake increases lactate production in hypoxia. **a**, Schematic of *BSG* KD CMs with glucose-derived pyruvate redirected toward lactate production with the addition of the mitochondrial pyruvate carrier (MPC) inhibitor UK-5099. **b,c** ^13^C-Glucose tracer contribution to pyruvate m+3 (**b**) and lactate m+3 (**c**) ± 250 uM UK-5099 in *BSG* KD CMs. Data are presented as mean ± SD. *n* = 6 replicates. Significance was assessed for **b** and **c** on the m+3 labeled metabolites, with a two-way ANOVA with Tukey’s multiple comparisons post hoc test, relative to the control: *BSG* KD vehicle control 21% O2.

**Figure S8.**
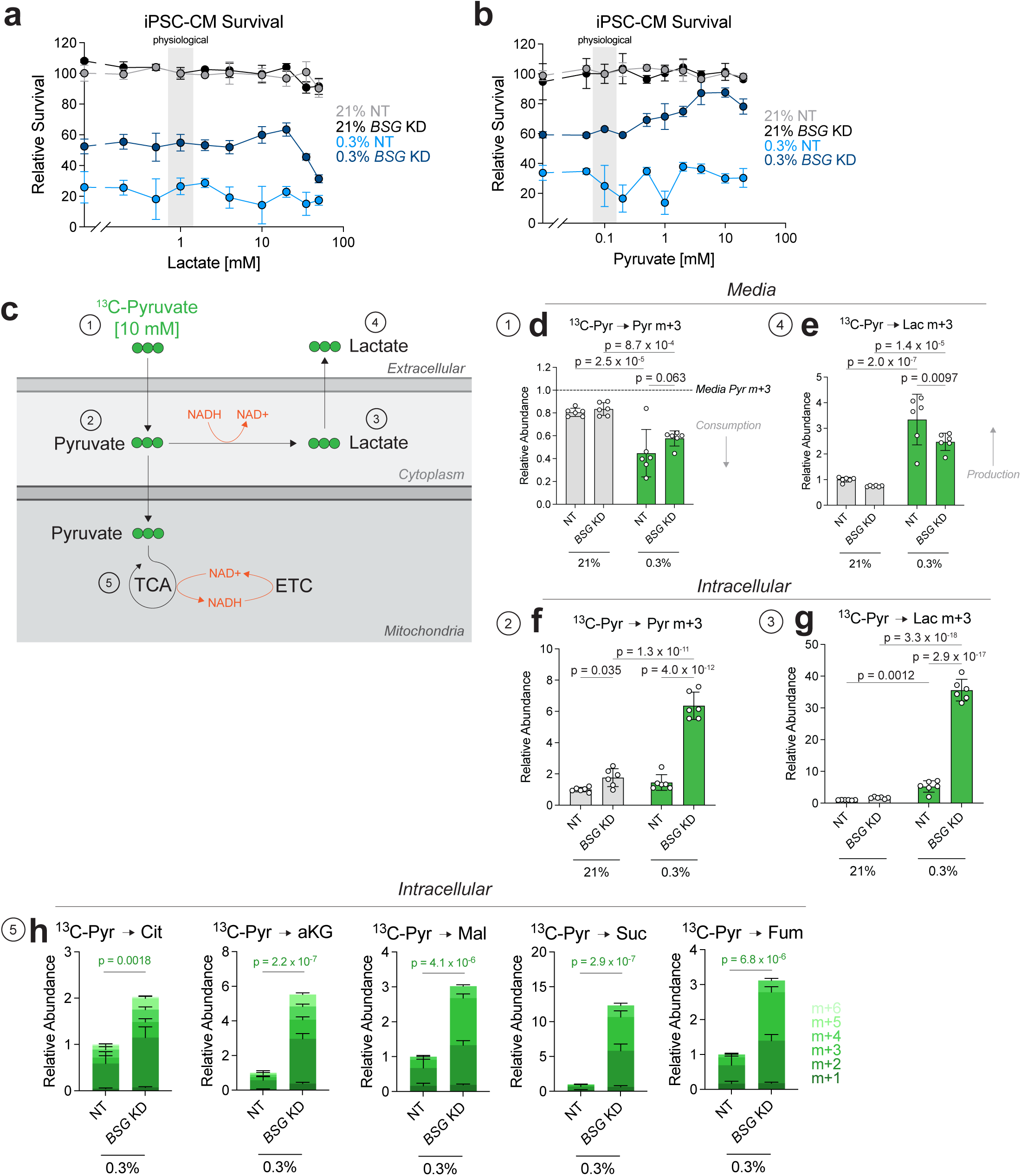
Pyruvate supplementation further rescues *BSG* KD but not NT CMs in hypoxia. **a,** Electron donor lactate dose escalation in NT and *BSG* KD CMs for 3 days in 21% O2 and 0.3% O2. The physiological concentration of lactate (1 mM) is highlighted. *n* = 3 replicates. **b,** Electron acceptor pyruvate dose escalation in NT and *BSG* KD CMs for 3 days in 21% O2 and 0.3% O2. The physiological concentration of pyruvate (0.1 mM) is highlighted. *n* = 3 replicates. **c,** NT and *BSG* KD CMs were treated with uniformly-labeled ^13^C-pyruvate tracer at a high dose (10 mM) for 24 hr in 21% O2 or 0.3% O2. Pyruvate can have two fates: (1) reduction to lactate to regenerate NAD+, or (2) oxidation in the mitochondria. Green circles represent ^13^C-labeled carbons. **d,** Proportion of ^13^C-pyruvate (m+3) remaining in media after 24 hr of culture with NT or *BSG* KD CMs in 21% O2 and 0.3% O2. Normalized to the amount of pyruvate m+3 detected in unconditioned media. *n* = 6 replicates. **e,** Labeled lactate m+3 production in the media, normalized to NT CMs in 21% O2. *n* = 6 replicates. **f**, Intracellular levels of pyruvate m+3, normalized to NT CMs in 21% O2. *n* = 6 replicates. **g,** Intracellular levels of lactate m+3, normalized to NT CMs in 21% O2. *n* = 6 replicates. **h,** Intracellular levels of labeled TCA cycle intermediates, normalized to NT CMs in 0.3% O2. Detected TCA cycle intermediates include, citrate, alpha-ketoglutarate, malate, succinate, and fumarate. *n* = 6 replicates. Data are presented as mean ± SD. Significance was assessed for **d-g** with a two-way ANOVA with Tukey’s multiple comparisons post hoc test, relative to the control: NT 21% O2, and for **h** with an unpaired two-tailed Student’s t-test comparing the sums of all the labeled isotopologues for each TCA cycle intermediate, relative to the control: NT 0.3% O2.

**Figure S9.**
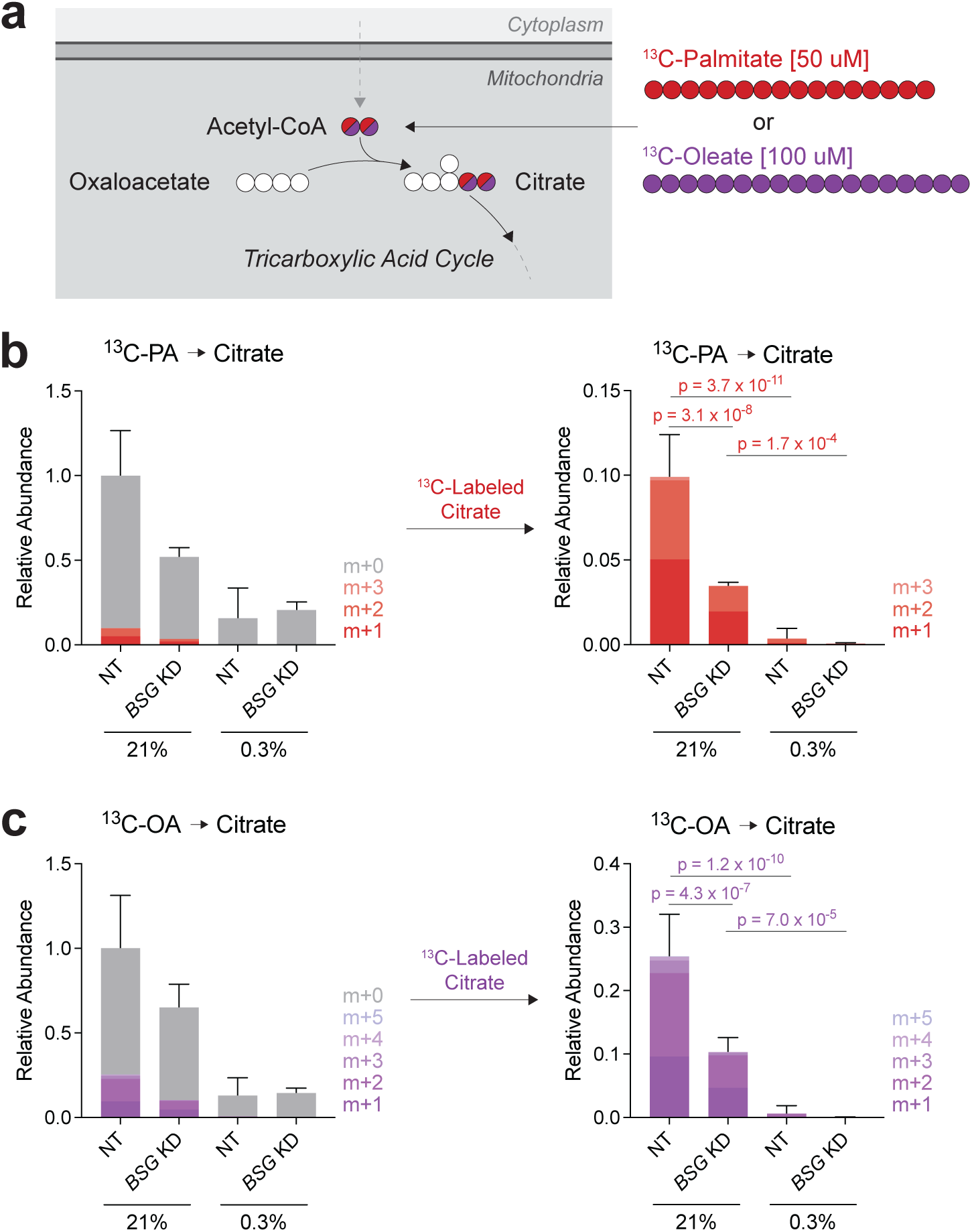
*BSG* KD decreases fatty acid contribution to the TCA. **a,** Schematic of ^13^C-palmitate or ^13^C-oleate contribution to the TCA. These labeled fatty acids were conjugated to BSA and given to iPSC-CMs for 24 hours of labeling after 2 days of culture in 21% O2 or 0.3% O2. Red and purple circles represent ^13^C-labeled carbons from ^13^C-palmitate and ^13^C-oleate, respectively. **b**, Relative abundance of total citrate, highlighting contribution from palmitate. *Left:* Data are presented as mean ± SD of total citrate. *Right:* ^13^C-labeled citrate from palmitate. *n* = 6 replicates. **c**, Relative abundance of total citrate, highlighting contribution from oleate. *Left:* Data are presented as mean ± SD of total citrate. *Right:* ^13^C-labeled citrate from oleate. *n* = 6 replicates. Significance was assessed for **b** and **c** comparing the sum of labeled citrate isotopologues with a two-way ANOVA with Tukey’s multiple comparisons post hoc test, relative to the control: NT 21% O2.

## METHODS

### Human iPSC culture and iPSC-cardiomyocyte differentiation

iPSCs were maintained in mTeSR Plus (Stem Cell Technologies 100-0276) and passaged with accutase (Innovative Cell Technologies AT104-500) and 10 μM ROCK Inhibitor Y-27632 onto matrigel (Corning 35623) coated flasks. iPSCs were cultured at 37 °C in a 5% CO2/95% air incubator. When iPSCs reached 80% confluency, mTeSR Plus was changed for RPMI 1640 medium supplemented with B27 supplement minus insulin and 12 μM CHIR-99021 on Day 0 for 24 hours to initiate iPSC-CM differentiation. iPSCs were differentiated into cardiomyocytes using a modified Wnt modulation protocol according to Lian et al. 2013, supplemented with 50 μg/mL Ascorbic Acid for the first 7 days and 0.5 μM SB-431542 (TGFb inhibitor) on days 2 and 3. On day 7, media was changed to RPMI 1640 medium with B27 supplement with insulin. iPSC-CMs were replated for lactate purification on day 15 using 15 U/mL papain (Worthington Biochemical LK003178) in HBSS to dissociate CMs. iPSC-CMs were purified with lactate purification media for 5 days according to Tohyama et al. 2013. Finally, lactate-purified iPSC-CMs were matured for 10 days by replacing standard media with a physiological media (RPMI 1640 no glucose with 2 mM glutamine, 4 mM glucose, 100 μM pyruvate, 1 mM L-lactate, 100 μM oleic acid BSA-conjugated, 50 μM palmitic acid BSA-conjugated, and 120 μM carnitine). Fatty acids were conjugated to BSA according to Knight et al. 2021.^17^

### Hypoxia culture of iPSC-CMs and iPSCs

For hypoxia culture, iPSC-CMs or iPSCs were moved to Billups-Rotherberg Modular Incubator Chambers (MIC-101) and flushed with a mixture of 0.3% O2, and 95% N2 / 5% CO2 for 2 minutes and then sealed. Modular chambers were incubated in a 37 °C incubator. Media was changed daily and the chambers were re-flushed with a fresh gas mixture. For moderate hypoxia experiments (5% O2), iPSC-CMs were moved to a standard incubator with 5% O2 and 5% CO2. For normoxia experiments (21% O2), iPSC-CMs or iPSCs were maintained in a standard incubator with 21% O2 and 5% CO2.

### iPSC-CM CRISPRi screen

Karyotype-normal WTC-11 iPSCs with CRISPRi dCas9-KRAB in the CLYBL locus (generous gift from Dr. Ken Nakamura) were expanded for a lentiviral infection with a CRISPRi library. We used a compact genome-wide library from Replogle et al, 2020, *Nature Biotechnology*, with two guide RNAs (gRNAs) per plasmid, both targeting the same gene. To produce lentivirus, plasmid library was packaged with HEK239T-17 cells using the *Trans*IT-Lenti Transfection Kit (Mirus Bio LLC MIR 6600). Filtered and concentrated virus was transduced into iPSCs using an MOI titration curve to target an MOI of 0.3. To maintain a minimum of 500X coverage in the screen, 37.5 million iPSCs were infected with lentivirus and 8 ug/mL polybrene during iPSC replating. A small subset of iPSCs were collected for flow cytometry to validate the percent infection during the screen. iPSCs were purified with 1.2 μg/mL puromycin for 5 days and subsequently seeded onto 11 T75 flasks for iPSC-CM differentiation. iPSC-CMs were lactate purified and a small subset was fixed for immunofluorescent labeling and flow cytometry to validate successful differentiation and expression of the gRNA construct. iPSC-CMs were matured with 10 days of culture in physiological media before moving into three O2 tensions (21%, 5%, 0.3% O2) for 9 days of culture with daily media changes. iPSC-CM survival was quantified by counting cells using trypan blue and a hemocytometer, where each T75 flask from the screen served as a replica.

Cell pellets were collected from each condition from the two replicates of the screen, and genomic DNA was isolated with the NucleoSpin Blood L Kit (Macherey-Nagel 740954.100). gRNA constructs were amplified with PCR using the universal forward primer (CAAGCAGAAGACGGCATACGAGATGCGGCCGGCTGTTTCCAGCTTAGCTCTTAAA) and with the following barcoded i5 reverse primers.

#### CRISPR screen PCR amplification barcoded primers

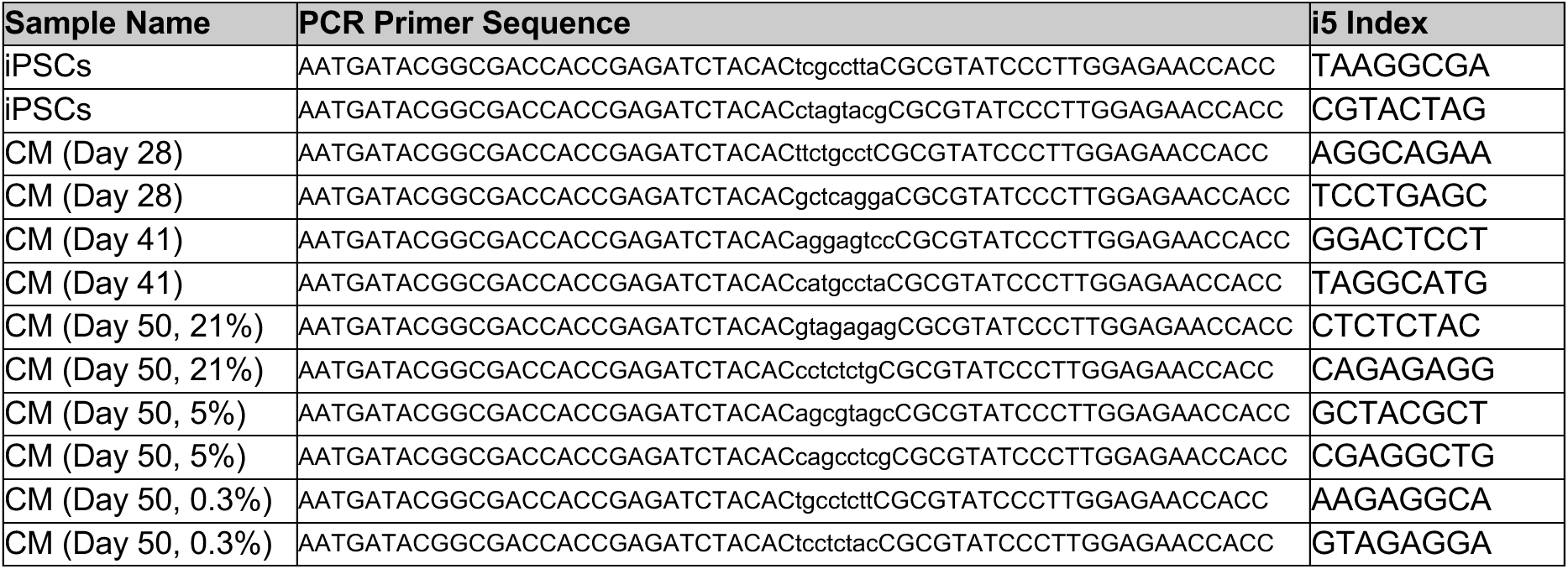

PCR amplification was performed with NEBNext Ultra II Q5 MasterMix (NEB M0544X) with the following cycling conditions: 98 °C for 30 sec, 22 cycles of [98 °C for 10 sec, 63 °C for 75 sec], followed by 72 °C for 5 min. We performed a 0.5-0.65X SPRI cleanup with PCR cleanup beads from the UC Berkeley Sequencing Facility. Samples were validated with a bioanalyzer, quantified with a Qubit, and pooled for paired-end sequencing onto a Novaseq 6000 with the following sequencing primers.

#### CRISPR screen sequencing primers

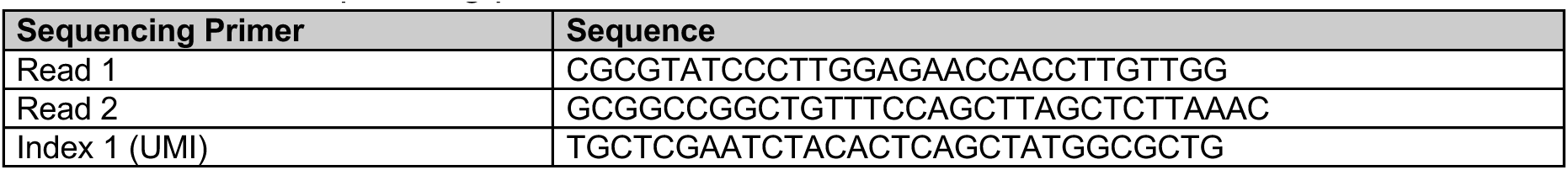

Two independent differentiations and screens were performed. Raw counts were median normalized by barcoded sample, and replicate 1 and 2 of the screens were averaged for each sample. Generally essential gRNAs (counts <15 in iPSCs) were filtered out, and iPSC-CM gRNAs that were essential for differentiation (counts <15 in iPSC-CMs Day 50 21% O2) were filtered out for both replicates of the screen. Log2FC comparisons were taken across different treatment groups, and values reported are the average of both screen replicates. GO Enrichment Analysis was performed with PANTHER software, using the top 200 essential gRNAs. Gene Set Enrichment Analysis (GSEA) was performed with 1,000 permutations for all known Complex IV subunits and assembly factors using the following gene list.

#### Complex IV subunits and assembly factors

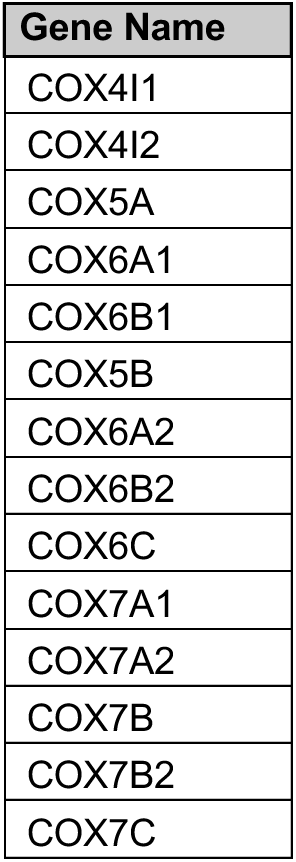
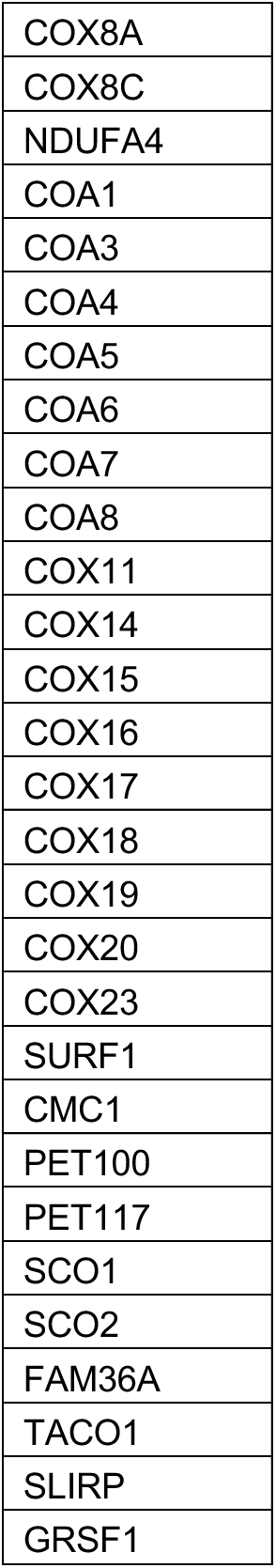

| Gene Name |
| --- |
| COX4I1 |
| COX4I2 |
| COX5A |
| COX6A1 |
| COX6B1 |
| COX5B |
| COX6A2 |
| COX6B2 |
| COX6C |
| COX7A1 |
| COX7A2 |
| COX7B |
| COX7B2 |
| COX7C |
| COX8A |
| COX8C |
| NDUFA4 |
| COA1 |
| COA3 |
| COA4 |
| COA5 |
| COA6 |
| COA7 |
| COA8 |
| COX11 |
| COX14 |
| COX15 |
| COX16 |
| COX17 |
| COX18 |
| COX19 |
| COX20 |
| COX23 |
| SURF1 |
| CMC1 |
| PET100 |
| PET117 |
| SCO1 |
| SCO2 |
| FAM36A |
| TACO1 |
| SLIRP |
| GRSF1 |

### CRISPRi Knockdowns in iPSC-CMs

To generate KDs in iPSC-CMs, iPSCs expressing dCas9-KRAB were transduced with lentivirus containing gRNA plasmids adapted from Replogle et al, 2020, *Nature Biotechnology*. Expression plasmids contained two gRNAs targeting the same gene, driven by human U6 and mouse U6 promoters. The plasmids additionally harbored puromycin resistance and BFP for visualization. Lentivirus was generated using psPAX2 and VSV-G packaging plasmids and X-tremeGENE transfection reagent (Sigma-Aldrich 6366236001) in HEK293T/17 cells. Virus was precipitated with Alstem Viral Precipitation Solution (VC125) and subsequently concentrated before transduction of iPSCs during replating. After 48 hr of infection, 1.2 μg/mL puromycin was added to the media for 5 days. Selected iPSCs were differentiated to iPSC-CMs. gRNA pairs used for KDs are included below.

#### CRISPRi Guide RNAs

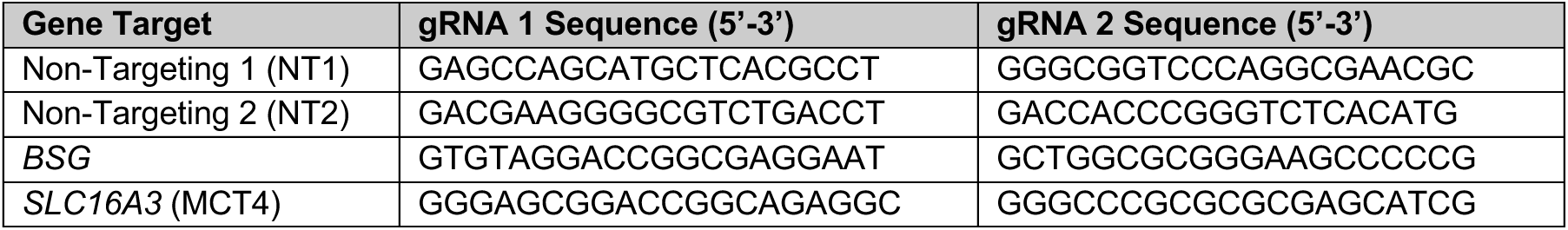

### Plasmid Construction

Dual gRNA plasmids were generated using the pJR103 backbone (Addgene #187242) which was a generous gift from Dr. Joseph Replogle. The backbone was PCR amplified with KOD Xtreme Hot Start DNA Polymerase (Sigma 71975), and dual gRNA cassettes were synthesized by TWIST and were inserted via Gibson Assembly (NEB E2621).

### Immunocytochemistry

Adherent iPSC-CMs were washed with PBS and fixed in 4% paraformaldehyde (Thermo 043368.9M) in PBS for 20 mins. Cells were permeabilized for 30 min in blocking buffer [0.05% Tween-20 (Promega H5152) in 0.5% BSA (Sigma A9418) in PBS] with 0.5% Triton X-100 (Sigma 9002-93-1), then blocked for 1 hr in blocking buffer with 1% normal donkey serum (Jackson Immuno Research Labs NC9624464). iPSC-CMs were co-stained with anti-cTnT (1:400) (Abcam, ab45932) and anti-Basigin (1:400) (R&D; AF972) for 1 hr at room temperature. iPSC-CMs were washed 3X in PBS and then incubated for 30 mins at room temperature in Hoechst (1:10,000) (Invitrogen 62249) and secondary antibodies: Alexa Fluor 488 anti-rabbit (1:400) (Invitrogen A21206) and Alexa Fluor 647 anti-goat (1:400) (Invitrogen, A-21447). iPSC-CMs were washed 3X in PBS and imaged with a Zeiss LSM880 Confocal Microscope.

### Flow Cytometry

Lactate-purified iPSC-CMs were dissociated with papain and fixed in 4% paraformaldehyde (Thermo 043368.9M) in PBS for 20 mins. Cells were permeabilized for 30 min in blocking buffer [0.05% Tween-20 (Promega H5152) in 0.5% BSA (Sigma A9418) in PBS] with 0.5% Triton X-100 (Sigma 9002-93-1), then blocked for 1 hr in blocking buffer with 1% normal donkey serum (Jackson Immuno Research Labs NC9624464). iPSC-CMs were co-stained for markers cTnT (1:400) (Abcam, ab45932) and ɑ-Actinin (1:400) (Sigma, A7811, (Clone EA-53)) for 1 hr at room temperature. Isotype control samples were stained with either Rabbit IgG (2.5 μg/mL) (Invitrogen, 10500C) or Mouse IgG1 (25 μg/mL) (R&D Systems, MAB002, Clone 11711). Cells were washed 3X in PBS and then incubated for 30 mins at room temperature in Hoechst (1:10,000) (Invitrogen 62249) and secondary antibodies: Alexa Fluor 647 anti-mouse IgG (1:400) (Invitrogen, A21202), and Alexa Fluor 555 anti-rabbit IgG (1:400) (Invitrogen, A31572). Cells were washed 3X in PBS and then resuspended in blocking buffer. Cells were analyzed on an Attune NxT Flow Cytometer for a minimum of 10,000 events per sample. Debris was gated out with SSC v FSC, doublets were gated out with FSC-Height v FSC-Area, and cells were included by Hoechst+ staining (both mono- and bi-nucleated cells). The percent purity of iPSC-CMs was determined by cTnT+/ɑ-Actinin double positivity, and the fraction of GFP+ (gRNA expressing) iPSC-CMs was determined by endogenous GFP signal over a no-virus control iPSC-CM line.

### Inhibitor Survival Experiments

iPSC-CMs were plated onto 96-well plates and were allowed to recover for at least 4 days before initiating experiments. Inhibitors were dissolved in DMSO and added to cell culture media up to 0.1% DMSO. Media with DMSO and drugs was changed on iPSC-CMs every 24 hr during survival experiments in 21% O2 and 0.3% O2. The following inhibitors were used in this study.

#### Inhibitors for dose response survival curves in iPSC-CMs

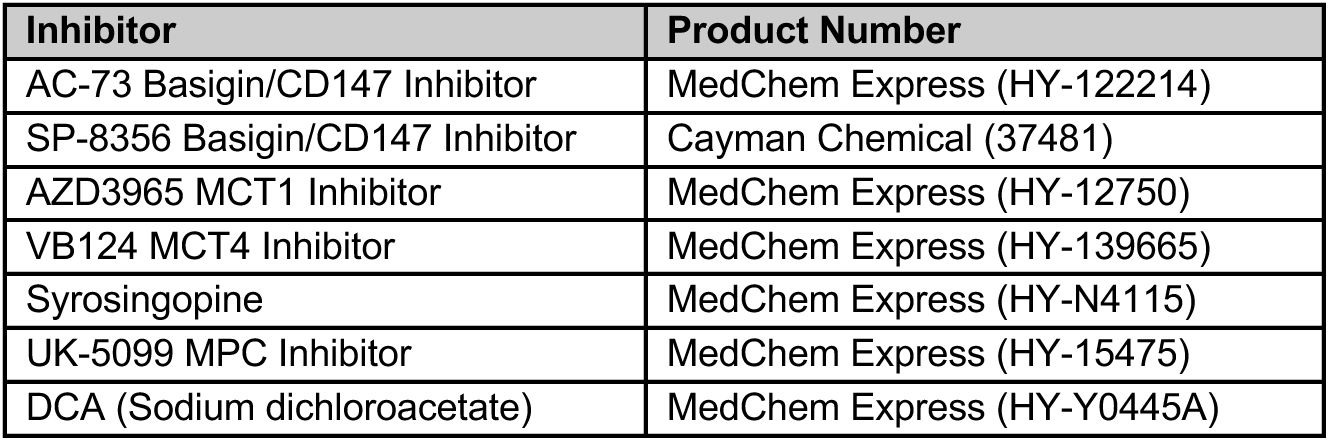

| Inhibitor | Product Number |
| --- | --- |
| AC-73 Basigin/CD147 Inhibitor | MedChem Express (HY-122214) |
| SP-8356 Basigin/CD147 Inhibitor | Cayman Chemical (37481) |
| AZD3965 MCT1 Inhibitor | MedChem Express (HY-12750) |
| VB124 MCT4 Inhibitor | MedChem Express (HY-139665) |
| Syrosingopine | MedChem Express (HY-N4115) |
| UK-5099 MPC Inhibitor | MedChem Express (HY-15475) |
| DCA (Sodium dichloroacetate) | MedChem Express (HY-Y0445A) |

### Western Blotting

For cell samples, cells were lysed in RIPA buffer (Thermo PI89901) with 1:100 Protease and Phosphatase Inhibitor Cocktail (78440) and 1:300 Benzonase Nuclease. For mouse heart samples, tissue was lysed with a Qiagen TissueLyser, using T-PER (Thermo 78510) with 1:100 Protease and Phosphatase Inhibitor Cocktail and 1:300 Benzonase Nuclease. Protein concentration was determined with the Rapid Gold BCA kit (Thermo A53225). 10 or 20 μg of protein was loaded per lane onto polyacrylamide gels (Bio-Rad 4561106) with Precision Plus Protein ladder (Biorad 1610374). Gels were run at 200V for 30 mins and transferred onto PVDF membranes (Biorad 1704157). Membranes were blocked in TBST with 5% milk for 1 hour on a rocker at room temp. Primary antibodies were diluted in 5% milk in TBST at 1:1000 and were incubated on the membranes overnight on a rocker at 4 °C. Membranes were washed 3X with TBST and secondary antibodies were diluted in 5% milk in TBST at 1:4000 and incubated for 1 hr on a rocker at room temp. Membranes were washed 3X with TBST and briefly treated with ECL substrate (Pierce 32106) before imaging on an Azure 600 Imager. The following antibodies were used in this study.

#### Information for primary antibodies for immunofluorescence (IF) and western blots (WB)

| Target | Dilution | Catalog Number |
| --- | --- | --- |
| Anti-α-Actinin | 1:400 (IF) | Sigma; A7811 (Clone EA-53) |
| Anti-Cardiac Troponin T | 1:400 (IF) | Abcam; ab45932 |
| Anti-Basigin | 1:400 (IF), 1:1000 (WB) | R&D; AF972 |
| Anti-Basigin | 1:1000 (WB) | R&D; AF772 |
| Anti-Vinculin | 1:1000 (WB) | Abcam; ab129002 |
| Anti-MCT1 | 1:1000 (WB) | Abcam; ab85021 |
| Anti-MCT4 | 1:1000 (WB) | Proteintech; 22787-1-AP |
| Anti-PDHA1 | 1:1000 (WB) | Invitrogen; 459400 |
| Anti-PDH-E1α (pSer232) | 1:1000 (WB) | Sigma-Aldrich; AP1063 |
| Anti-PDH-E1α (pSer293) | 1:1000 (WB) | Cell Signaling; 37115 |
| Anti-PDH-E1α (pSer300) | 1:1000 (WB) | Sigma-Aldrich; AP1064 |
| Anti-PDK1 | 1:1000 (WB) | Proteintech; 18262-1-AP |
| Anti-PDK2 | 1:1000 (WB) | Proteintech; 15647-1-AP |
| Anti-PDK3 | 1:1000 (WB) | Abcam; ab322408 |
| Anti-PDK4 | 1:1000 (WB) | Proteintech; 12949-1-AP |

### Proteomics

After 3 days of culture in 21% O2 and 0.3% O2, iPSC-CMs were lifted with papain, pelleted, and stored at −80 °C. Cell pellets were lysed using the Lysis Solution from the EasyPep MS Sample Prep Kit (Thermo). Protein quantity was determined with the Rapid Gold BCA kit (Thermo), and 100 μg of protein was reduced and alkylated. Protein was digested with Trypsin/Lys-C Protease Mix with shaking at 37 °C for 3 hr. Finally, peptides were cleaned-up and sent to UC-Davis Proteomics core to dry the peptide sample and resuspended in 0.1% formic acid in water for LC-MS analysis. Peptide samples were run on the Exploris480 mass spectrometer. Samples were median normalized and the reported protein levels are relative to the NT 21% O2 condition. Two independent iPSC-CM differentiations and hypoxia/normoxia exposures were collected with *n* = 3 replicates each. One differentiation is presented as a representative.

### RNA Isolation

iPSC-CMs or iPSCs in 24-well plates were lysed with RLT Buffer (Qiagen 79216) with 1% β-mercaptoethanol and moved to an eppendorf tube. Total RNA was extracted using the RNeasy Mini Kit (Qiagen). All RNA samples were treated with DNAse to remove potential DNA contamination. The quantity and purity of RNA was determined by nanodrop.

### RT-qPCR

Complementary DNA (cDNA) was synthesized with the iScript cDNA Synthesis Kit (Bio-Rad 1708891). For the PCR reaction, cDNA was combined with Maxima SYBR Green Kit (Thermo K0222) and amplified on the Applied Biosystems QuantStudio 5 Real-Time PCR System for 95 °C for 10 min followed by 40 cycles of 95°C for 15 sec, 60 °C for 30 sec, and 72 °C for 30 sec.

Each cell treatment had a minimum of triplicates, and average cycle threshold (Ct) values were averaged across two technical replicates for each cDNA sample. Expression levels were normalized to *GAPDH* or *ACTB* expression using the ΔCt method, and fold change values across conditions were determined with the ΔΔCt method. All primers were ordered from Integrated DNA Technologies and are listed below.

#### Sequences of human primers used for qPCR

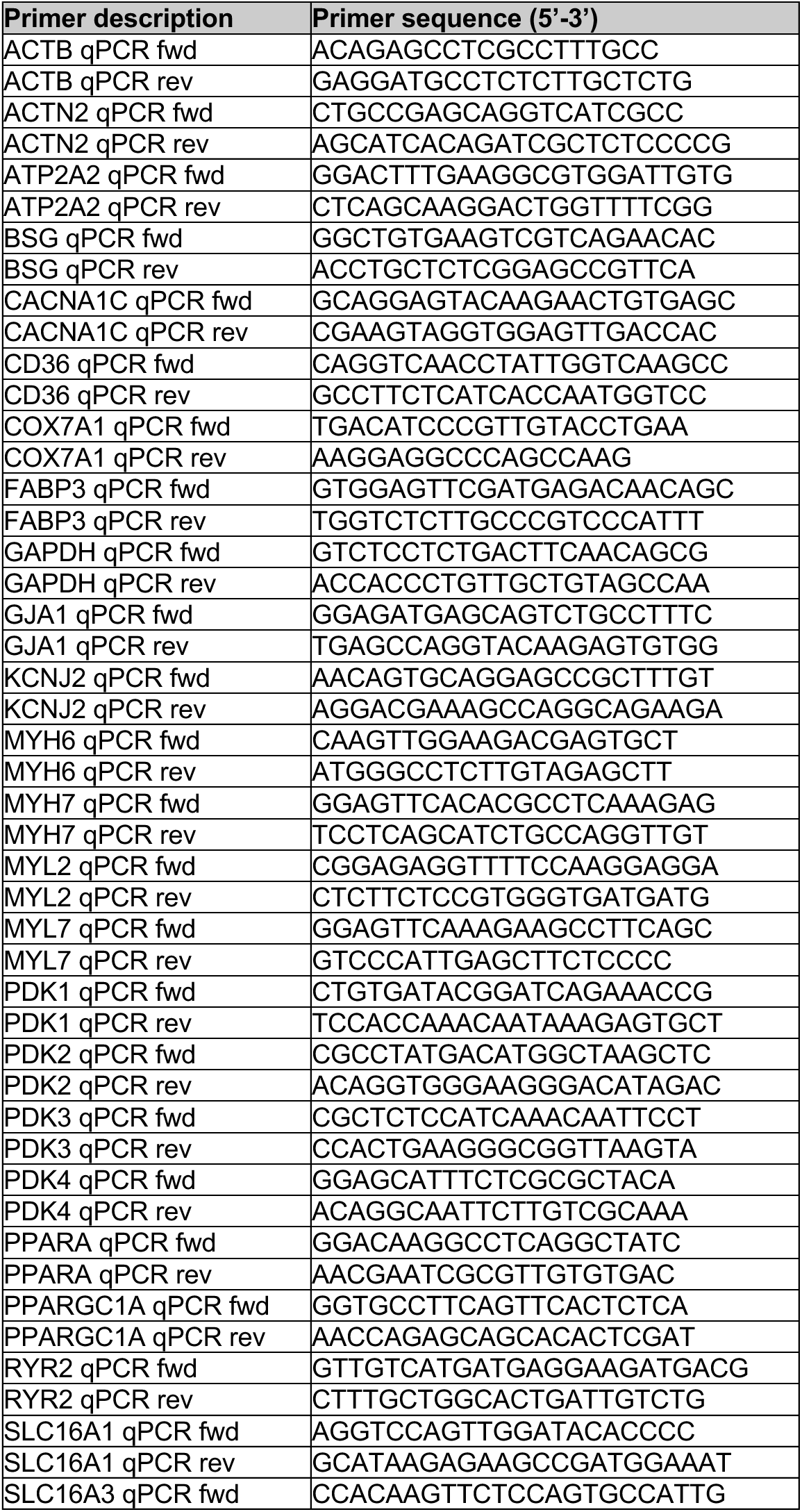
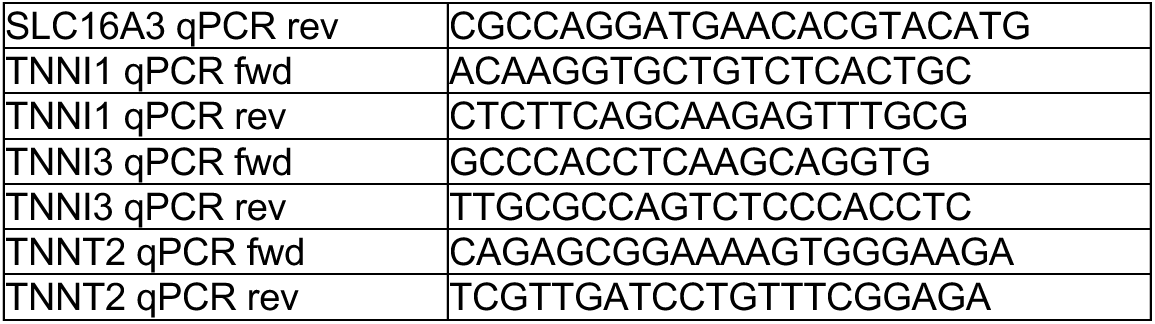

### iPSC-CM polar metabolite extraction

2.85 x 10^5^ iPSC-CMs were plated on matrigel-coated 24-well plates and were allowed to recover for at least 4 days before beginning hypoxic treatments. Physiological media was changed daily and after 3 days of treatment, culture media was removed and cells were washed with PBS. Polar metabolites were extracted by quenching the cells with 300 μL of 80% Methanol with 100 μM D8-valine internal standard. Plates were incubated at −80 °C for 30 mins. Cells were scraped and transferred to eppendorf tubes, briefly vortexed, and incubated on ice for 20 mins. Samples were centrifuged at 20,000 g at 4 °C for 20 mins. Supernatant was moved to new tubes and was lyophilized overnight. The remaining protein pellet was quantified with a BCA for sample normalization. Lyophilized samples were resuspended in 60% acetonitrile and sonicated for 3 mins. Samples were placed on a rotator at 4 °C for 20 mins, then spun down at 20,000 g at 4 °C for 20 mins. Supernatant was moved to glass autosampler vials for LC-MS analysis.

### Conditioned media polar metabolite extraction

iPSC-CMs in 24-well plates received daily media changes with physiological media. After 2 days of culture in 21% O2 or 0.3% O2, fresh physiological media was replaced and iPSC-CMs were placed back into their respective O2 conditions. 24 hr later, 100 μL conditioned media was collected from each well. Media was centrifuged at 350 g for 5 mins to pellet cell debris. 50 μL of supernatant was moved to a new tube and polar metabolites were extracted by adding 450 μL of 80% Methanol with 100 μM D8-valine internal standard. Samples were vortexed, placed at −80 °C for 30 minutes, then spun down at 20,000 g at 4 °C for 10 mins. Supernatant was moved to a new tube and samples were lyophilized and resuspended as described above. To determine metabolite enrichment or depletion from baseline media, no cell (media alone) controls were extracted (*n* = 6 wells).

### Stable isotope labeled tracer experiments in iPSC-CMs

Stable isotope metabolic tracers
| Tracer | Product Number | Concentration of Tracer |
| --- | --- | --- |
| D-glucose (U-<sup>13</sup>C<sub>6</sub>, 99%) | Cambridge Isotope Laboratories Inc. (CLM-1396-2) | 4 mM |
| Sodium L-lactate (3-<sup>13</sup>C, 98%) | Cambridge Isotope Laboratories Inc. (CLM-1578-0.25) | 1 mM |
| Palmitic acid (U-<sup>13</sup>C<sub>16</sub>, 98%) | Cambridge Isotope Laboratories Inc. (CLM-409-0.5) | 50 μM |
| Oleic acid (U-<sup>13</sup>C<sub>18</sub>, 98%) | Cambridge Isotope Laboratories Inc. (CLM-460-0.005) | 100 μM |
| Sodium-pyruvate (U-<sup>13</sup>C<sub>3</sub>, 99%) | Cambridge Isotope Laboratories Inc. (CLM-2440-0.1) | 10 mM |

Stable isotope-labeled media was prepared using RPMI 1640 without glucose (Gibco 11879020) base media. Each missing component of the physiological media was added to the base media, and various tracers were used instead of their unlabeled counterparts at the following concentrations. After 2 days of culture in 21% O2 or 0.3% O2, media was removed and cells were washed with PBS before adding fresh media with tracers. Cells were placed back into their respective O2 conditions. 24 hours later, polar metabolites were extracted from media and cells separately. The following metabolic tracers were used in this study.

#### Stable isotope metabolic tracers

To quantify the abundance of different metabolite isotopologues, Thermo Fisher TraceFinder software was used to verify raw peak areas. These distributions were corrected for natural isotope abundance using AccuCor open-source code^67^. These corrected peak areas were normalized to D8-valine internal standard peak area, and then to a BCA to account for differences in cell survival across conditions.

### Liquid chromatography-mass spectrometry

Samples were analyzed on an Orbitrap Exploris 240 high-resolution mass spectrometer (Thermo Fisher Scientific) equipped with an electrospray ionization source. The instrument was interfaced with a Vanquish Horizon ultra-high-performance liquid chromatography system operating under hydrophilic interaction chromatography conditions. For each sample, 2 μL of the polar metabolite extract was injected onto an iHILIC-(P) Classic column (2.1 × 150 mm, 5 µm; HILICON AB, 160.152.0520). The column temperature was maintained at 40 °C, while the autosampler temperature was kept at 4 °C. Mobile phase A consisted of 20 mM ammonium bicarbonate in water, adjusted to pH 9.6 using ammonium hydroxide, and mobile phase B was acetonitrile. Chromatographic separation was performed at 200 μL/min using the following gradient: 0–18 min, 85% to 20% B; 20–20.5 min, linear increase from 20% to 85% B; 20.5–28 min, held at 85% B. Each run was preceded by a 10-minute equilibration at initial conditions.

The mass spectrometer was run in full-scan polarity-switching mode with spray voltages of 3.5 kV (positive) and 3.25 kV (negative). Data were collected from m/z 70–1000 at a resolution setting of 60,000. Instrument parameters included an RF Lens level of 60%, an AGC target of 1e7, and a maximum injection time of 200 ms. Gas settings were as follows: sheath gas 35 units, auxiliary gas 10 units, and sweep gas 0.5 units. The ion transfer tube temperature was set to 300 °C, and the vaporizer temperature to 35 °C.

### Semi-targeted metabolomics

For semi-targeted metabolomic analyses, the Mass Spectrometry Metabolite Library of Standards (MSMLS; IROA Technologies) was run on the LC–MS platform. Fragmentation data were generated by analyzing the standards in both positive and negative ionization modes using data-dependent MS2 (ddMS2) acquisition. Higher-energy collisional dissociation (HCD) collision energies of 15%, 45%, and 90% were applied, and MS2 spectra were collected on the Orbitrap at a resolution of 30,000. To support compound identification, pooled quality control (QC) samples were generated by mixing equal volumes of each experimental sample. These QC samples were analyzed under the same ddMS2 conditions using a targeted mass list derived from detectable MSMLS metabolites. Untargeted compound identification was also performed with AcquireX, using the same ddMS2 acquisition parameters.

Compound Discoverer (Thermo Fisher Scientific) was used for compound identification and peak integration, referencing both a custom spectral library generated from the MSMLS standards and the mzCloud online spectral database.

### Animal handling

Mice were maintained under standard housing conditions with controlled temperature and humidity (∼20 °C and ∼50% relative humidity) on a 12-hour light/12-hour dark cycle. Animals were provided standard laboratory chow (PicoLab 5058) ad libitum. All animal procedures were conducted in accordance with protocols approved by the Institutional Animal Care and Use Committee at UCSF and the Gladstone Institutes.

To generate a cardiomyocyte-specific conditional *Bsg* knockout (KO) mouse model, *Bsg^fl/fl^*;*Myh6*-Cre mice (hereafter referred to as *Bsg^CKO^*) were used. *Bsg^fl/fl^* mice were generously provided by Dr. Romana A. Nowak (University of Illinois, Urbana-Champaign), and *Myh6*-Cre mice (The Jackson Laboratory, stock no. 011038) were generously provided by Dr. Arun Padmanabhan (University of California, San Francisco). Mice were bred such that all offspring were homozygous for *Bsg^fl/fl^*, with either the presence (*Myh6*-Cre^+^; *Bsg^CKO^*) or absence (*Myh6*-Cre^-^; floxed control) of the *Myh6*-Cre transgene. Pups were weaned at 3-4 weeks of age, and mice used for heart collection were 6-9 months old. Mice were anesthetized with 3% isoflurane in oxygen, sacrificed by cervical dislocation, and hearts were rapidly excised and flash-frozen in liquid nitrogen.

### NADH/NAD assay

The whole-cell NADH/NAD+ ratio was measured using the NAD/NADH Glo Assay from Promega (G9072). Briefly, iPSC-CMs were cultured in 21% or 0.3% O2 for 3 days. Cells were lysed and prepared according to manufacturer instructions. Luminescence was measured with a SpectraMax plate reader.

